# Allosteric regulation and crystallographic fragment screening of SARS-CoV-2 NSP15 endoribonuclease

**DOI:** 10.1101/2022.09.26.509485

**Authors:** Andre Schutzer Godoy, Aline Minalli Nakamura, Alice Douangamath, Yun Song, Gabriela Dias Noske, Victor Oliveira Gawriljuk, Rafaela Sachetto Fernandes, Humberto D. Muniz Pereira, Ketllyn Irene Zagato Oliveira, Daren Fearon, Alexandre Dias, Tobias Krojer, Michael Fairhead, Alisa Powell, Louise Dunnet, Jose Brandao-Neto, Rachael Skyner, Rod Chalk, Frank von Delft, Dávid Bajusz, Miklós Bege, Anikó Borbás, György Miklós Keserű, Glaucius Oliva

## Abstract

SARS-CoV-2 is the causative agent of COVID-19. The highly conserved viral NSP15 endoribonuclease, NendoU, is a key enzyme involved in viral immune evasion, and a promising target for the development of new classes of antivirals. Yet, the broad variety of recognition sequences, complex assembly and kinetics, and lack of structural complexes hampers the development of new competitive or allosteric inhibitors for this target. Here, we performed enzymatic characterization of NendoU in its monomeric and hexameric form, showing that hexamers are allosteric enzymes with a positive cooperative index of 2. By using cryo-EM at distinct pH’s combined with X-ray crystallography and structural analysis, we demonstrate the potential for NendoU to shift between open and closed states, and assembly in larger supramolecular entities, which might serve as a mechanism of self-regulation. Further, we report results from a large fragment screening campaign against NendoU, revealing multiple new allosteric sites for the development of inhibitors.

## Introduction

The current coronavirus disease 2019 (COVID-19) pandemic, caused by Severe Acute Respiratory Syndrome Coronavirus 2 (SARS-CoV-2), has become a major global health and economic crisis (*1, 2*). The rapid and global spread of COVID-19 has resulted in an urgent need for the development of effective therapeutic options against this novel coronavirus (*3*). SARS-CoV-2 is a member of the *Betacoronavirus* genus, which includes SARS-CoV and MERS-CoV (*1, 4*). Its genome of ∼ 30 kb consists of a long replicase gene encoding non-structural proteins (NSPs), followed by structural and accessory genes (*5*). Due to a ribosomal frameshifting, the replicase gene encodes two open reading frames (ORFs), rep1a and rep1b, that are translated into two large polyproteins, pp1a and pp1ab (*6*). The cleavage of these results in sixteen NSPs with distinct functions, including several replicative enzymes which are key targets for drug development (*7*).

One of the least understood NSPs of coronaviruses is NSP15, a highly conserved nidoviral RNA uridylate- specific endoribonuclease (NendoU), carrying the catalytic domain of the endonuclease family (*8, 9*). NendoU is three-domain 34 kDa protein that was demonstrated to form a barrel shaped hexamers in solution by the assembly of two trimers (*8, 10*). The oligomerization is mostly driven by the contacts of the N-terminal (ND) and the middle (MD) domains, while the C-terminal (CD) domain carry the canonical catalytic residues from the endonuclease family (*11*). In the oligomer, the six active sites seem to be independently located at the top and bottom of the complex. Despite this, NendoU shows maximum activity as an oligomer (*12*), the exact role of the hexamer is unknown.

Due its unique activity, as well as the fact that this enzyme was co-localized with the *de-novo* synthetized viral RNA (*13*) and the viral RNA dependent RNA polymerase (NSP12) (*14*), a key protein of the viral replication machinery (*15*), suggesting that NendoU is directly involved in the viral RNA metabolism. Moreover, evidence suggests that NendoU is crucial for coronaviruses innate immune evasion due its antagonistic IFN effects, promoted by the decreasing host cell dsRNA sensors, which was observed in macrophages and *in vivo* (*16–20*). Still, the exact role of NendoU in viral metabolism remains unclear, but accumulated evidence point that NendoU is a key viral enzyme and therefore a valuable target for antiviral development or even attenuated viral vaccines (*17, 21*). Still, the complex RNA binding mode of NendoU, lack of structural complexes and allosteric behavior hamper the development of new antivirals.

Here, we combine X-ray crystallography, cryo-electron microscopy (cryo-EM) and biochemical analysis to investigate the complex biochemical profile of SARS-CoV-2 NendoU in both monomeric (NendoU^mon^) and hexameric (NendoU^hex^) forms. Combined structural and biochemical analysis indicate that NendoU activity depends on the hexamer dynamic and suggest that supramolecular interactions are involved in the enzyme regulatory mechanism. Furthermore, we carried out a large-scale X-ray fragment screen to probe new putative druggable sites. Our data sheds light in the understanding of NendoU enzymatic profiles, opening the path for the development of new allosteric and competitive inhibitors.

## Materials and Methods

### Protein production and purification

The SARS-COV-2 cDNA sample was gently donated by Dr. Edilson Durigon (GenBank MT126808.1), which was generated using SCRIPT One-Step RT-PCR (Cellco Biotec) and random hexamer primers. NSP15 coding sequence was amplified using FastPol (Cellco Biotec) with primers 5’-CAGGGCGCCATGAGTTTAGAAAATGTGGCTTTTAATG-3’ and 5’-GACCCGACGCGGTTATTGTAATTTTGGGTAAAATGTTTCTAC-3’, and inserted pETM11/LIC (gently donated by Dr. Arie Geerlof) using Ligation-Independent Cloning method (*22*). Final construct pETM11-NSP15 included a N-terminal 6xHis-tag followed by a TEV cleaving site and residues 1-346 of NSP15.

Transformed BL21(DE3) cells were growth in LB Lennox medium supplemented with kanamycin at 37°C until DO_600_ = 1.0 and keep it for 16h at 18°C after induction with 0.5 mM isopropyl β-D-1-thiogalactopyranoside. Harvested cells were resuspended in Buffer A (50 mM TRIS-HCl pH 8.0, 300 mM NaCl, 10% glycerol, 20 mM Imidazole) supplemented with 0.1 mg.mL^−1^ lysozyme, 10 U of benzonase (Cellco Biotec) and 1.0 mM DTT, and disrupted using sonication. Cleared lysate was passed through a HisTrap HP 5 mL (GE Healthcare) equilibrated in Buffer A, and them protein was eluted with Buffer A supplemented with 300 mM imidazole. Excessive imidazole was removed using a Sephadex XK 26/60 column (GE Healthcare) equilibrated with Buffer A. For obtaining NSP15 in monomeric form (NendoU^mon^), sample was incubated overnight at 8°C with 4 mM DTT and 0.1 mg.mL^−1^ TEV protease, than passed through a HisTrap HP 5 mL equilibrated in Buffer A to remove non-cleaved protein and TEV protease. For obtaining NSP15 in hexameric form (NendoU^hex^), 6xHis-tag was not cleaved. For both forms, final purification was performed by size exclusion chromatography using a HiLoad Superdex 75 16/60 (GE Healthcare) equilibrated with 20 mM HEPES pH 7.5, 150 mM NaCl, 5% (v/v) glycerol, 0.5 mM TCEP. Purity was confirmed by SDS-PAGE 12%. Protein concentration was estimated using the theoretical extinction coefficient at 280 nm of 33,140 M^−1^.cm^−1^ (*23*). Mutations were generated by inverse-PCR method.

### Endoribonuclease assay and kinetic analysis

Fluorogenic oligonucleotide substrate 5’6-FAM-dArUdAdA-6-TAMRA3’ was purchased from GenScript. Unless state otherwise, activity assays were performed in 25 mM BIS-Tris-HCl buffer 6.0 pH, 0.25 μM substrate and 100 nM NendoU^mon^ or 25 nM NendoU^hex^ at 35°C. Time-curse reactions were directly monitored in a Stratagene Mx3005P (Agilent Technologies) using FAM filters. initial velocity (v_0_) was estimated using the slope from linear regressions of reaction monitored during minutes 3-10. Analysis and graphs were performed with OriginPro v9.0.

The gel-based RNA cleavage assay was performed to verify the cleavage of the PolyU biotinylated RNA substrate 5’-UGACCUCUAGCUAGAGGUCA(U)_30_-3’ by nsp15, based on (*24*). A 15% acrylamide/urea gel (8 M) was pre-runned at 200 V for 30 min in TBE buffer. Samples in 1X formamide loading buffer (Gel loading buffer II – Ambion) were heated for 5 min at 95 ºC prior to analysis. The denaturing gel was run for 2 h at 200 V and RNA bands were visualized with SYBR Safe DNA Gel Stain (Invitrogen) under UV light.

### CryoEM sample preparation and data collection

For sample preparation, 3 µl of 0.5 mg/ml NendoU^hex^ in experiment buffer were applied on the Quantifoil 300 mesh copper R1.2/1.3 grid. Buffers tested were 50 mM Tris-HCl pH 7.5 mM, 200 mM NaCl or 50 mM BIS-Tris HCl pH 6.0, 200 mM NaCl or 50 mM PBS pH 6.0, 200 mM NaCl. Samples containing oligoDT were made in 50 mM PBS pH 6.0, 200 mM NaCl buffer containing with 5 µM of oligoDT_20_ (Thermofisher™). The sample was blotted for 4 s (4 C, 100% humidity) with a blot force of -12 and then plunge frozen using a Vitrobot Mark IV (Thermo Fisher). Frozen samples were imaging on Titan Krios (G2) microscope operated at 300 kV. Movies were collected on a K3 detector in super resolution counting mode with a slit width of 20 eV and at a nominal magnification of 105kx corresponding to a calibrated physical pixel size of 0.831 Å at the specimen level. Data acquisition was done using EPU software (version 2.6) with a defocus range of 1.4 - 3.2 nm. Data collections statistics is summarized in Table S2.

### CryoEM data processing and modelling

In summary, all movies were aligned using MotionCor2 (*25*), and processed using cryoSPARC v2.15 (*26*). Contrast transfer function (CTF) parameters for each micrograph were determined by Patch CTF. Initial templates were construct using manual picking tools, and them all micrographs were picked using Template Picker, followed by Extraction with box size of 256 px and two rounds of 2D classification. Selected particles were used for Ab-initio modelling and two rounds of Heterogeneous Refinement. Selected volumes were refined using Non-uniform Refinement and D3 symmetry. Detailed data processing flowchart scheme for samples in buffers 50 mM Tris-HCl pH 7.5 mM, 200 mM NaCl or 50 mM BIS-Tris HCl pH 6.0, 200 mM NaCl or 50 mM PBS pH 6.0, 200 mM NaCl are available in Figs. S6, S7 and S8, respectively, and final gold standard FSC resolution for the models were respectively 2.98, 3.2 and 2.5 Å. For modelling, ChimeraX was used for rigid fitting of model using PDB 7KF4 as template, followed by real space refinement using Phenix (*27, 28*). Data collections and model refinement statistics are summarized in Table S2. To analyze variability of SARS-CoV-2 NSP15 structural flexibility, we used the 3D Variability Analysis (3DVA) tool available in cryoSPARC v2.15 (*29*). Particles and mask were used to compute 3 eigenvectors of the covariance matrix of the data distribution using 3DVA (*29*), with resolution filtered at 4 Å. Model series were generated using 3DVA display tool. Movies were generated with ChimeraX v1.2.5.

### Helical processing for NSP15 filaments

Helix processing was performed with relion v3.1 (*30*). Data was aligned with MotionCorrv2 (*31*), and CTF was corrected with ctffind v4.1.13 (*32*). Particle picking was performed manually with relion Manual helical picking tool using 120 Å particle diameter. Particle were extracted (2437) with box of 600 pixels as helical segments (tube diameter 150 Å and start-end mode, helical rise 10, number of asymmetry units 6). Particles were classified with Class2D as Helix segments, with 150 Å tube diameter, bimodal angular search, angular search psi 6 degrees and helical rise 10 Å.

### Crystallization, data collection and data processing for of NendoU X-ray models

Crystals of NendoU^mon^ were obtained in multiple conditions. For that, protein was concentrated to 3.4 mg/mL and crystallized in sitting drop at 18°C by vapor diffusion. Hexagonal crystals containing a dihedral asymmetric unit were obtained in 15% PEG 8000, 0.1 M Sodium/Potassium Phosphate pH 6.2 and 20 % w/v Polyethylene glycol 3350, 100 mM BIS-TRIS propane, pH 6.5, 200 mM Sodium sulfate, and cryo-condition was obtained by adding 20% (v/v) ethylene glycol. Orthorombic crystals containing hexagonal asymmetric unit were obtained with protein incubated with 100 µM oligoDT and crystallized in 0.1 M trisodium citrate pH 5, 14 % w/v PEG6000, and cryo-condition was obtained by adding 20% (v/v) ethylene glycol.

Data was processed with XDS via autoPROC (*33, 34*), scaled with Aimless via CCP4(*35*), solved by molecular replacement using PDB 6×1b as template via Phaser (*36*), and refined with phenix.refine or BUSTER (*37, 38*). Model was built with Coot and validated with Molprobity (*39, 40*) Fig.s were made using PyMOL, ChimeraX and APBS (*28, 41*). Data collection and refinement statistics are available in Table S3. Conservation analyses were performed with Consurf (*42*), and results are available in Fig S14.

### Fragment screening of SARS-CoV-2 NendoU

For the fragment screening campaign, protein crystals were formed using 300 nL of NendoU^mon^ at 3.4 mg/mL mixed with 300 nL of well solution containing 14% (w/v) PEG 6,000 and 0.1 M tri-sodium citrate pH 5.0 and 10 nL of seeds stocks of the same condition in Swissci 96-Well 3-Drop plates (Molecular Dimensions) containing 30 µL of well solution. Crystals were formed after 4 days at 20 °C by vapor diffusion. For soaking, 40 nL of each fragment compound from XCHEM Poised Library (*43*) and OPEN-EU DRIVE fragment library (final concentration of 100 mM) were added to a crystallization drop using an ECHO acoustic liquid handler dispenser at the Diamond light source XChem facility (*44*) Cryo-condition was created by adding 20% ethylene glycol to the drops using ECHO acoustic liquid handler. Crystals were soaked for two hours before being harvested using XChem SHIFTER technology, and data was collected at the beamline I04-1 in automated mode. The XChem Explorer pipeline (*45*) was used for structure solution with parallel molecular replacement using DIMPLE (*46*). Fragments were identified using PANDDA software (*47*) and CLUSTER4X (*48*). Data was modelled and refined using phenix.refine, BUSTER and COOT (*37–39*), and validated using Molprobity (*40*). Coordinates and structure factors of ground-state model was deposited in PDB under the code 5SBF. Bound-state statistics and PDB codes are summarized in Supplementary table 1.

### Native electrospray mass spectrometry (ESI-TOF) intact mass analysis

Proteins for native mass spectrometry were held on ice and buffer exchanged into 75 µl of 50 mM ammonium acetate pH 7.5 by 3 rounds of gel filtration using BioGel P6 (Biorad) spin columns according to the manufacturer’s instructions. Mass spectra were acquired using an Agilent 6530 QTOF operating in positive ion 1 GHz mode using a standard ESI source. Samples were introduced via a syringe pump at a flow rate of 6 µl/min. The ion source was operated with the capillary voltage at 3500 V, nebulizer pressure at 17 psig, drying gas at 325°C and drying gas flow rate at 5 L/min. The instrument ion optic voltages were as follows: fragmentor 430 V, skimmer 65 V and octopole RF 750 V. M/z spectra were analysed using Masshunter B.07.00 (Agilent); ESIprot (*49*) and using an ion table.

### Chemical synthesis of nucleoside analogs

5’-azido-5’-deoxy-thymidine (LIZA-7) was prepared according to the procedure we published recently.(*50*) (The first synthesis of this compound was reported in 2007.(*51*)) 5’-Deoxy-5’-thiothymidine (FUZS-5) was prepared according to a novel procedure reported here, through the deacetylation of 5’-S-Acetyl-5’-deoxy-5’-thiothymidine, with the latter being synthesized based on the procedure reported by Kawai et al.(*52*) (The first synthesis of 5’-Deoxy-5’-thiothymidine via the deacetylation of the 3’,5’-diacetyl analog was reported in 1964.(*53*)) Reagents were purchased from Sigma-Aldrich Chemical Co. and used without further purification. Optical rotations were measured at room temperature with a Perkin-Elmer 241 automatic polarimeter. TLC was performed on Kieselgel 60 F_254_ (Merck) with detection by UV-light (254 nm) and immersing into sulfuric acidic ammonium molybdate solution or 5% ethanolic sulfuric acid followed by heating. Flash column chromatography was performed on silica gel 60 (Merck, 0.040-0.063 mm). The ^1^H NMR (400 MHz) and ^13^C NMR (100 MHz) spectra were recorded with Bruker DRX-400 spectrometer at 25 °C. Chemical shifts are referenced to Me_4_Si (0.00 ppm for ^1^H) and to the residual solvent signals (CDCl_3_: 77.2, DMSO-d_6_: 39.5, CD_3_OD: 49.0 for ^13^C). MALDI-TOF MS analyses of the compounds were carried out in the positive reflectron mode using a BIFLEX III mass spectrometer (Bruker, Germany) equipped with delayed-ion extraction. 2,5-Dihydroxybenzoic acid (DHB) was used as matrix and F_3_CCOONa as cationising agent in DMF. ESI-TOF MS spectra were recorded by a microTOF-Q type QqTOFMS mass spectrometer (Bruker) in the positive ion mode using MeOH as the solvent.

#### 5’-S-Acetyl-5’-deoxy-5’-thiothymidine (Th-5’SAc, LIZA-7)

PPh_3_ (6.5 g, 24.6 mmol, 2.0 equiv.) was dissolved in dry THF (50 mL) and cooled to 0 °C. Diisopropyl azodicarboxylate (DIAD) (5.0 mL, 24.6 mmol, 2.0 equiv.) was added dropwise and stirred for 30 min at 0 °C. Thymidine (3.0 g, 12.3 mmol) and HSAc (1.8 mL, 24.6 mmol, 2.0 equiv.) were dissolved in dry DMF (50 mL) and added dropwise to the reaction mixture and stirred for 30 min at 0 °C and 30 min at r.t. The solvent was evaporated under reduced pressure and the crude product was purified by flash column chromatography (gradient elution, CH_2_Cl_2_/MeOH 97.5/2.5→95/5) to give **5’-S-Acetyl-5’-deoxy-5’-thiothymidine** (1.7 g, 46%) as a yellowish solid.

R_f_ = 0.21 (CH_2_Cl_2_/MeOH 95/5), [α]_D_ = +40.9 (*c* 0.11, DMSO),^1^H NMR (400 MHz, DMSO) *δ* (ppm) 7.45 (d, *J* = 0.8 Hz, 1H, H-6), 6.16 (dd, *J* = 7.6, 6.5 Hz, 1H, H-1’), 5.43 (d, *J* = 4.3 Hz, 1H, O*H*), 4.11 (td, *J* = 6.7, 3.4 Hz, 1H, H-3’), 3.80 – 3.71 (m, 1H, H-4’), 3.23 (dd, *J* = 13.8, 5.8 Hz, 1H, H-5’a), 3.11 (dd, *J* = 13.8, 7.3 Hz, 1H, H-5’b), 2.37 (s, 3H, AcC*H*_3_), 2.28 – 2.18 (m, 1H, H-2’a), 2.06 (ddd, *J* = 13.5, 6.2, 3.3 Hz, 1H, H-2’b), 1.81 (s, 3H, thymine C*H*_3_). ^13^C NMR (100 MHz, DMSO) *δ* (ppm) 194.9 (1C, Ac*C*O), 163.7, 150.5 (2C, C-2, C-4), 136.1 (1C, C-6), 109.9 (1C, C-5), 84.5, 83.9, 72.6 (3C, C-1’, C-3’, C-4’), 38.0 (1C, C-2’), 31.1 (1C, C-5’), 30.6 (1C, Ac*C*H_3_), 12.2 (1C, thymine *C*H_3_). MALDI-ToF MS: *m/z* calcd for C_12_H_16_N_2_NaO_5_S ^+^ [M+Na]^+^ 323.0672, found 323.0660.

#### 5’-Deoxy-5’-thiothymidine (FUZS-5)

5’-S-Acetyl-5’-deoxy-5’-thiothymidine (1.7 g, 5.7 mmol) was dissolved in dry MeOH (25 mL) under Ar, and NaOMe (458 mg, 8.5 mmol, 1.5 equiv.) was added and stirred at r.t. for 4 h. The reaction mixture was neutralized with Amberlite IR 120 H^+^ ion exchange resin, filtered, and evaporated under reduced pressure. The crude product was purified by flash column chromatography (CH_2_Cl_2_/MeOH 97.5/2.5→95/5) to give **5’-deoxy-5’-thiothymidine (FUZS-5)** (826 mg, 59%) as a white solid. R_f_ = 0.20 (CH_2_Cl_2_/MeOH 95/5), m.p. 200-202 °C; [α]_D_ = +5.0 (*c* 0.28, DMSO), ^1^H NMR (400 MHz, DMSO) *δ* (ppm) 7.51 (s, 1H, H-6), 6.19 (t, *J* = 6.4 Hz, 1H, H-1’), 5.36 (s, 1H), 4.22 (s, 1H), 3.78 (t, *J* = 5.9 Hz, 1H), 2.75 (t, *J* = 7.1 Hz, 1H), 2.49 (d, *J* = 15.6 Hz, 1H), 2.21 (dt, *J* = 13.7, 6.9 Hz, 1H), 2.06 (dd, *J* = 11.8, 4.4 Hz, 1H), 1.80 (s, 3H, thymine C*H*_3_). ^13^C NMR (100 MHz, DMSO) *δ* (ppm)163.7,150.5 (2C, C-2, C-4), 136.1 (1C, C-6), 109.9 (1C, C-5), 87.1 (1C, C-1’), 83.7 (1C, C-4’), 71.9 (1C, C-3’), 38.1 (1C, C-5’), 26.3 (1C, C-2’), 12.2 (1C, thymine *C*H_3_). MALDI-ToF MS: *m/z* calcd for C_10_H_14_N_2_NaO_4_S ^+^ [M+Na]^+^ 281.0566, found 281.0540.

## Results and Discussion

### Recombinant NendoU can be obtained as monomers or hexamers

NendoU^mon^ and NendoU^hex^ were purified as described. NendoU^mon^ was obtained after cleavage of the N-terminal histidine tag as a single monodisperse peak by gel filtration (Fig. S1). This diverges from previous reports, where NendoU was purified as a mix of monomers, trimers and hexamers using similar conditions (*12, 54*). However, all these authors also reported issues removing the N-terminal histidine tag. A similar gel filtration profile was obtained by retaining the N-terminal histidine tag, therefore suggesting that hexamerisation is induced by the extra N-terminal residues (Fig. S2). The importance of internal NSP15 N-terminal residues for proper oligomerization was already described for SARS-CoV NendoU, where the authors demonstrated that mutants such as E3A were exclusively expressed as monomers or trimers (*12*). Another possibility is that the N-terminal TEV site is only exposed in the monomeric form, while the hexamer offers some steric hindrance that blocks cleavage. Nevertheless, mass spectrometry confirms that monomers consist of cleaved protein while hexamers contain full-length protein (Fig. S3). Native mass spectra of NendoU^mon^ and NendoU^hex^ showed the former consists of folded monomer as the principal species with higher order species as minor components, while NendoU^hex^ consists of folded hexamer as the principal species with folded monomer and dodecamer (M12) as less abundant species (Fig. S4). Native mass spectra also indicate that uncleaved monomers have higher thermal stability than cleaved ones, suggesting stabilization promoted by the N-terminal tag (Fig. S4). The two samples obtained were enzymatically characterized and used in structural studies to help understand the function of oligomerization in NendoU activity and regulation.

### NendoU activity is controlled by pH

The activity of NendoU^mon^ and NendoU^hex^ were tested in multiple buffers, pH’s and salts concentrations, using a fluorogenic RNA oligo (*55*). For NendoU^mon^, significant activity was only observed in slightly acid buffers, such as sodium acetate pH 5.0 (v_0_ = 19.6 ± 0.12 RFU.min^−1^), sodium cacodylate pH 6.0, 150 mM NaCl (v_0_ = 15.5 ± 0.08 RFU.min^−1^), MES pH 6.0, 150 mM NaCl (v_0_ = 16.5 ± 0.07 RFU.min^−1^) and BIS-TRIS pH 6.5, 150 mM NaCl (v_0_ = 15.6 ± 0.14 RFU.min^−1^). A similar behavior was seen for NendoU^hex^, that presented maximum activity at sodium cacodylate pH 6.0, 150 mM NaCl (v_0_ = 998.4 ± 31.2 RFU.min^−1^) and BIS-TRIS pH 6.5, 150 mM NaCl (v_0_ = 902.8 ± 13.90 RFU.min^−1^). For more basic buffers, such as MOPS pH 7.0, HEPES pH 7.5 and TRIS pH 8.5, the enzyme activity of both NendoU^mon^ and NendoU^hex^ was considerably negligible. The overall reaction rate for NendoU^hex^ was much higher than NendoU^mon^ for all tested conditions. Still, this diverges from previous reports where hexamers were correlated with the enzymatic activity (*56, 57*). It was also possible to observe that NendoU^hex^ enzymatic activity is inversely proportional to salt concentrations, with a 3-fold increment of the activity in HEPES pH 7.5 in comparison with the same buffer added with 500 mM. Further studies were therefore conducted in 50 mM BIS-TRIS Buffer pH 6.0. All enzymatic activity values and detailed buffer are available in Table S1. Our data indicates that hexameric form is critical for enzyme activity, as monomeric form retains about 1-5% of enzymatic activity in similar conditions.

### NendoU is an allosteric enzyme with a positive cooperativity index

While a NendoU^mon^ time-course exhibited a classical first order reaction, NendoU^hex^ time-course clearly showed a positive-cooperative reaction, with a slow initial velocity that enters the exponential phase after 5 min (Fig. 1A). We therefore tested selected temperatures to determine the time took for NendoU^hex^ to reach half of the maximum activity at the exponential phase (t_1/2_). The t_1/2_ for temperatures of 25, 30, 35 and 40°C was 53.0, 29.7, 18.2 and 11.4 min, respectively (FIG. 1A). The data shows that higher temperatures not only influence the initial velocity, but also the positive cooperativity index of NendoU^hex^. We also observed a linear relationship between the 3-10 min interval for temperatures of 35 and 40°C, with R^2^ of 0.96 and 0.97, respectively. (Fig. 1A). Subsequently, all further analysis were performed at 35°C, and initial velocity (v_0_) was estimated using the slope from linear regressions of reaction monitored during 3-10 min. Using these conditions, we were able to calculate the values of *K*_0.5_ 3.9 ± 0.5 µM for NendoU^hex^, and *K*_m_ of 7.4 ± 1.0 µM for NendoU^mon^, indicating that the hexamer form has an increased affinity for the substrate (Fig. 1B and 1C). For NendoU^hex^, we also determined a Hill constant of 1.9 ± 0.4, indicating a positive cooperativity between the sites of NSP15 (Fig. 1B).

**Fig. 1.**
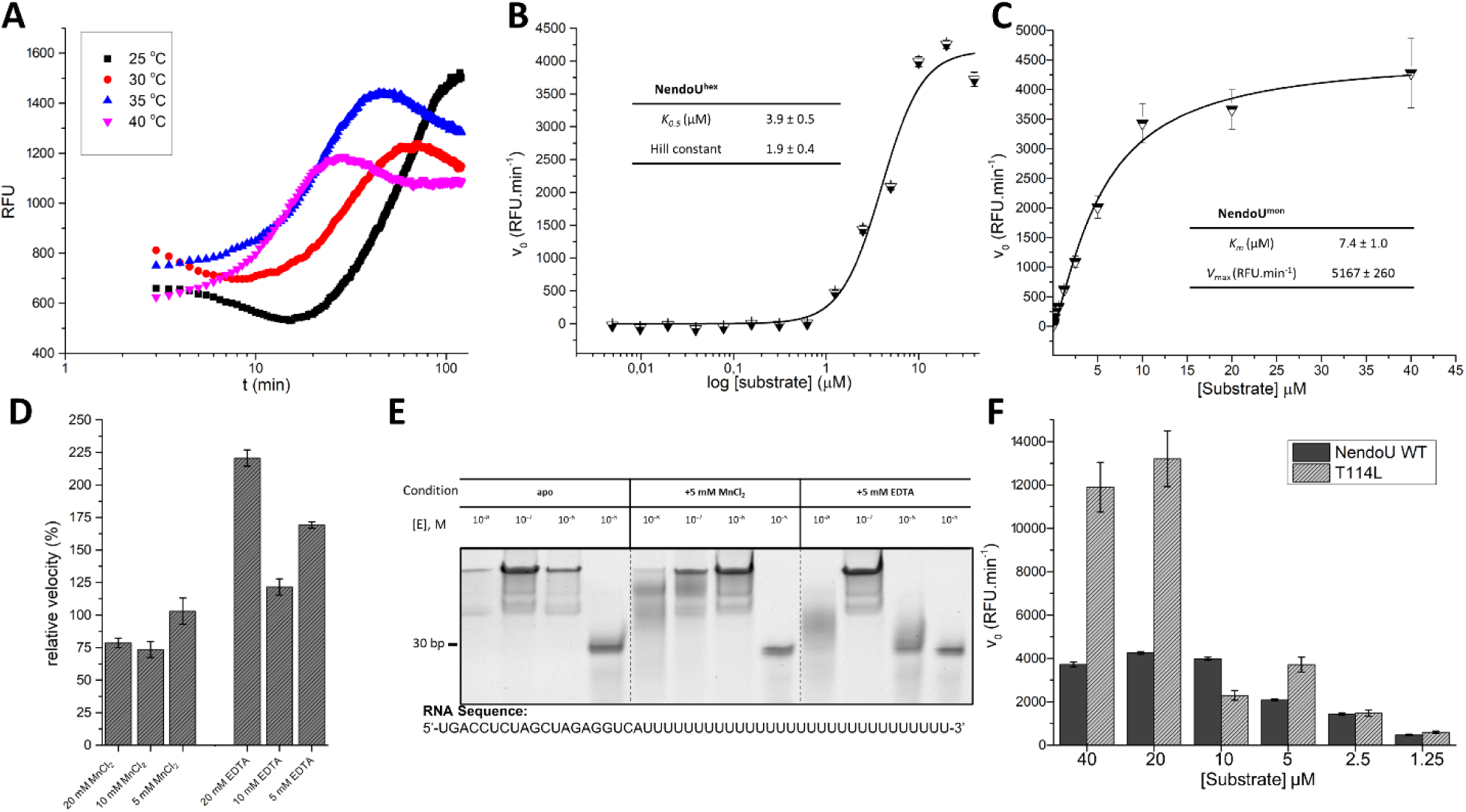
Summarized enzymatic profile characterization of NendoU. A) Time course reaction of NendoU^hex^ in different temperatures. B) Calculate Hill constant for NendoU^hex^, showing a positive cooperative index of 2. C) Michaelis Mentem plot of NendoU^mon^, showing a typical first order enzymatic profile. D) Enzymatic relative activity of NendoU^hex^ in presence of different concentrations of MnCl_2_ and EDTA. E) Acrilamide gel showing the activity of NendoU^hex^ against PolyU RNA in presence of different concentrations of MnCl_2_ and EDTA. F) Calculated initial velocities of mutant T114L and WT NendoU^hex^.

### NendoU is not manganese dependent

In terms of substrate specificity, the coronaviruses NendoU is described to cleave a great variety of single- and double-stranded RNAs in a Mn^2+^-dependent reaction, by hydrolyzing the 3’ end of pyrimidines, with a strong preference for uracil, and to release 2’, 3’-cyclic phosphate and 5’-hydroxyl ends as products (*11, 58, 59*). However, despite the many studies claiming the enhanced activity of NendoU in presence of Mn^2+^ (*7, 58, 60*), the ion was never observed to form complex with the polymer or substrates and products (*7, 58, 60*). Furthermore, the structural comparison of the NendoU active site with well characterized ion-independent endonucleases, like *Bos taurus* RNAse A, show minor differences between the position of key catalytic residues (*7, 11*), and the ion was also never observed to complex with the crystal’s structures of NendoU. Notwithstanding, during our structural studies described below, we collected multiple fluorescence spectra from different crystals, but neither of them indicated Mn in the samples (data not showed). Similarly, differential scanning fluorimetry showed no significant shift in enzyme stability in presence or absence of any Mn ions (data not shown).

We also tested the enzymatic activity of NendoU^hex^ in presence of several additives, including MnCl_2_ and MnSO_4_, and none of them seen to show a significant enhancement in the enzyme ratio. Next, we tested MnCl_2_ as an additive in various concentrations, and none of those seen to show to enhance the enzyme activity (Fig. 1D). Furthermore, we also tested the enzyme activity in high concentrations of EDTA (an obvious but missing control in most of above-mentioned studies) and the activity of NendoU^hex^ was not diminished in any of the tested concentrations, and even enhanced in some cases (Fig. 1D). To investigate its effect in a non-fluorescent assay, we performed an acrylamide/urea denaturating RNA page analysis in reactions containing NendoU^hex^ and a synthetic poly-U RNA oligo, for with or without Mn and using EDTA as control. Manganese was not observed to be crucial for the RNA cleavage, and EDTA does not seem to have any deleterious effect on the reaction (Fig. 1E). We therefore found evidences to claim that NendoU^hex^ is dependent of manganese or any other metal ions.

### NendoU binding to RNA involves active site and switch regions

At the middle surface of the barrel structure, we can find three pronounced cavities between each dimer subunits, which resembles the letter S (*56*) (Fig. 2A). Due to its pronounced size and proximity to the active site, it was assumed that this S cavity would serve to accommodate and recognize the variety of RNAs that can be processed by NendoU (*11, 56*). However, despite the many X-rays (*8, 54, 60, 61*) and Cryo-EM (*11*) structures of β-coronaviruses NendoU available to date, none of them show any evidence of a nucleic acid or nucleosides bound to the S cavity. Moreover, we observed that electrostatic potential calculations of NendoU suggests a strong negatively charged surface for this cavity (Fig. 2A), which is the opposite of what is expected for canonical DNA/RNA binding proteins recognition sites (*62*).

**Fig 2.**
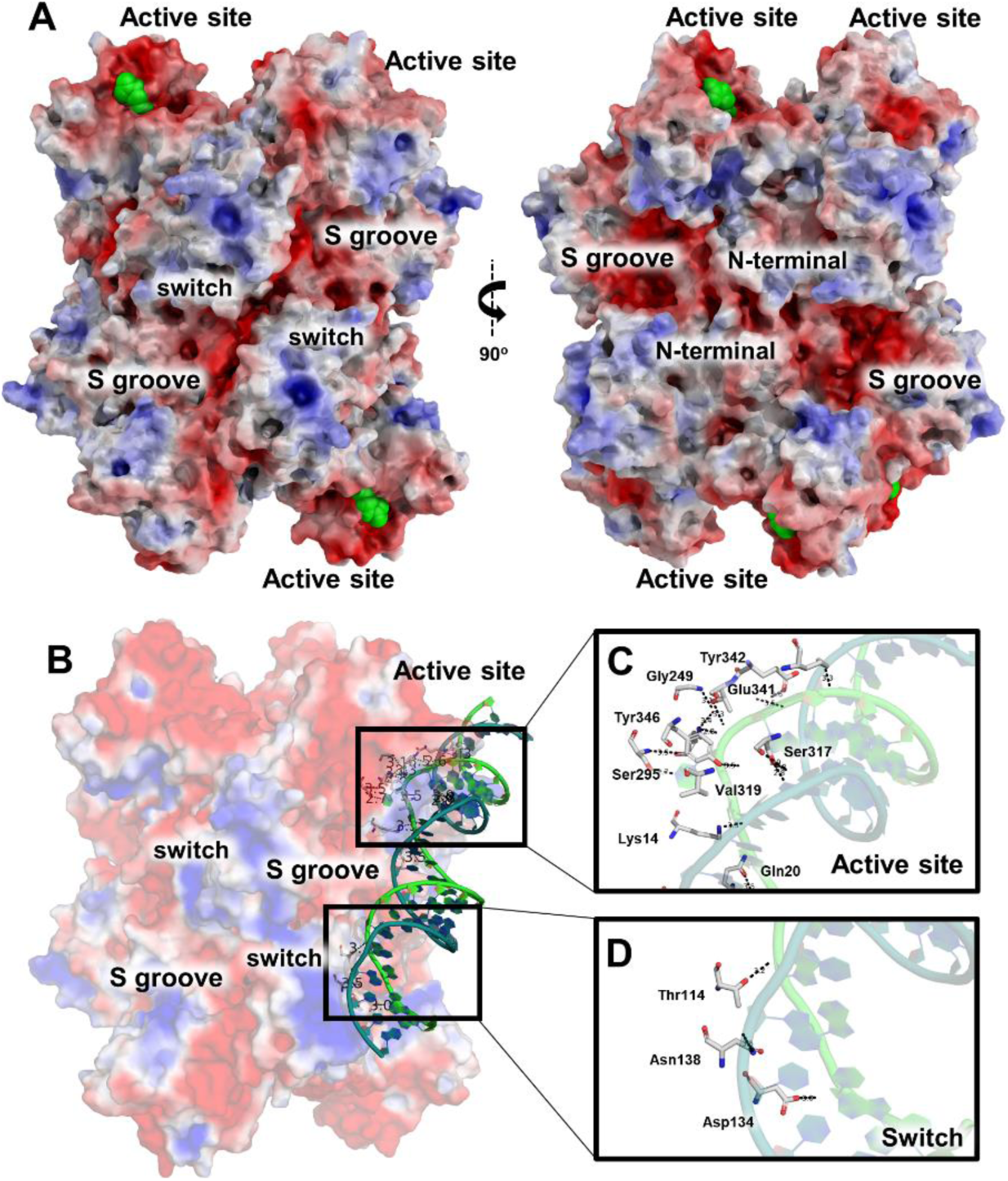
Overview of NendoU structure. A) Two distinct views of NendoU hexamer surface (PDB 7KF4), highlighting regions of interest, including the active site, the S groove and the switch region. Citrate molecule on the active site are depicted as green spheres. B) NendoU-dsRNA binding mode in the surface of the hexamer, occupying one active site and interacting with switch region. C) Detailed dsRNA binding mode on the active site of NendoU. D) Detailed dsRNA binding mode on the switch region of NendoU. Structure of NendoU is colored according to its electrostatic potential projected on surface charge (-5 to 5 f kJ/mol/e in red-white-blue color model). dsRNA is colored in green/blue and was depicted from PDB 7TJ2.

In fact, a recent cryo-EM study of NendoU with dsRNA shows that the RNA interacts not only with the active site, but also with the positively charged region (which we have named switch, an interface region comprising residues 96-121 from MD) from an adjacent nsp15 chain (Fig. 2B), but not with the S cavity (*63*). The structure revealed the binding mode of 5’ guanine π-π stacking with Trp332 (4.1 Å) at +2 subsite, and the 3’ processed uridine base bound at the +1 site, showing an unusual inversion of the base moiety angles to accommodate the interactions of its O2 with His249 NE2 (3.1 Å) and O4 with Leu345 N (4.5 Å). These interactions are key to understand not only the specificity of uridine site for P1 uridine, as well as its preferences for purine bases at the +2 site (*56, 60*). This could also be observed in our experiment using the poly-U synthetic RNA, where the cleavage shows that the 30-uracil tail is intact after reaction, highlighting the importance of a +2 purine for the proper cleavage (Fig. 1E).

The binding of the dsRNA to NendoU surface also revealed the interactions seem to be mediated by the positively charged residues of NendoU and the phosphate groups of the ribonucleosides (Fig. 2C and D). The lack of any major interactions between sugar and base moieties with the enzyme, as well as the flattened disposition of these subsites would explain the ability of NendoU in processing a broad variety of uridine containing sequences (*56, 57, 64*). At the negative subsites, which are located at the top edge of the NendoU barrel, we also notice the absence of any obvious cavity that would recognize any base in a specific manner, agreeing with the observation that multiple 3’ extensions can be recognized and processed by the enzyme (*56, 57, 64*). We can also see that the enzyme surface of this area is strong negatively charged, which would cause the rapid expelling of any nucleic acid products after cleavage (Fig. 2A). All this considered, the specificity of NendoU seems to be restricted to the uridine site at +1, and purine site at +2, narrowing our window for using structure-based methods for the development of competitive inhibitors.

### Cryo-EM reveals that NendoU switch conformation is pH-dependent and determines open-closed conformations

During our cryo-EM studies, we were able to elucidate multiple high-quality models of NendoU^hex^ in different pH’s and buffers, with resolution ranging from 2.5-3.5 Å (Fig. S6-S8). Despite its similarities, models showed consistent differences in the region of the switch (Fig. 3). For samples at HEPES pH 7.5, we observed that the canonical form of interacting switch regions assumes a more closed and mobile conformation in comparison with known crystal structures and cryo-EM models (Fig. 3A). However, in cryo-EM models obtained in PBS pH 6.0, we saw a less mobile and more open conformation of switch regions, much more like known crystal structures (Fig. 3B). The overall structure of the hexamer is also deeply affected by these distinct conformations of the switch region, where in pH 7.5 we observe a more contracted form of the barrel (Fig. 3A). The 3DVA Supplementary movies #1 and #2 show details of the contracting front and top view of the closed form of NendoU. In contrast, in pH 6.0 conditions, where enzyme is more active, we see that the structure is more rigidly defined in the open conformation, as can be seen in Supplementary movies #3 and #4 and in Fig. 3B.

**Fig 3.**
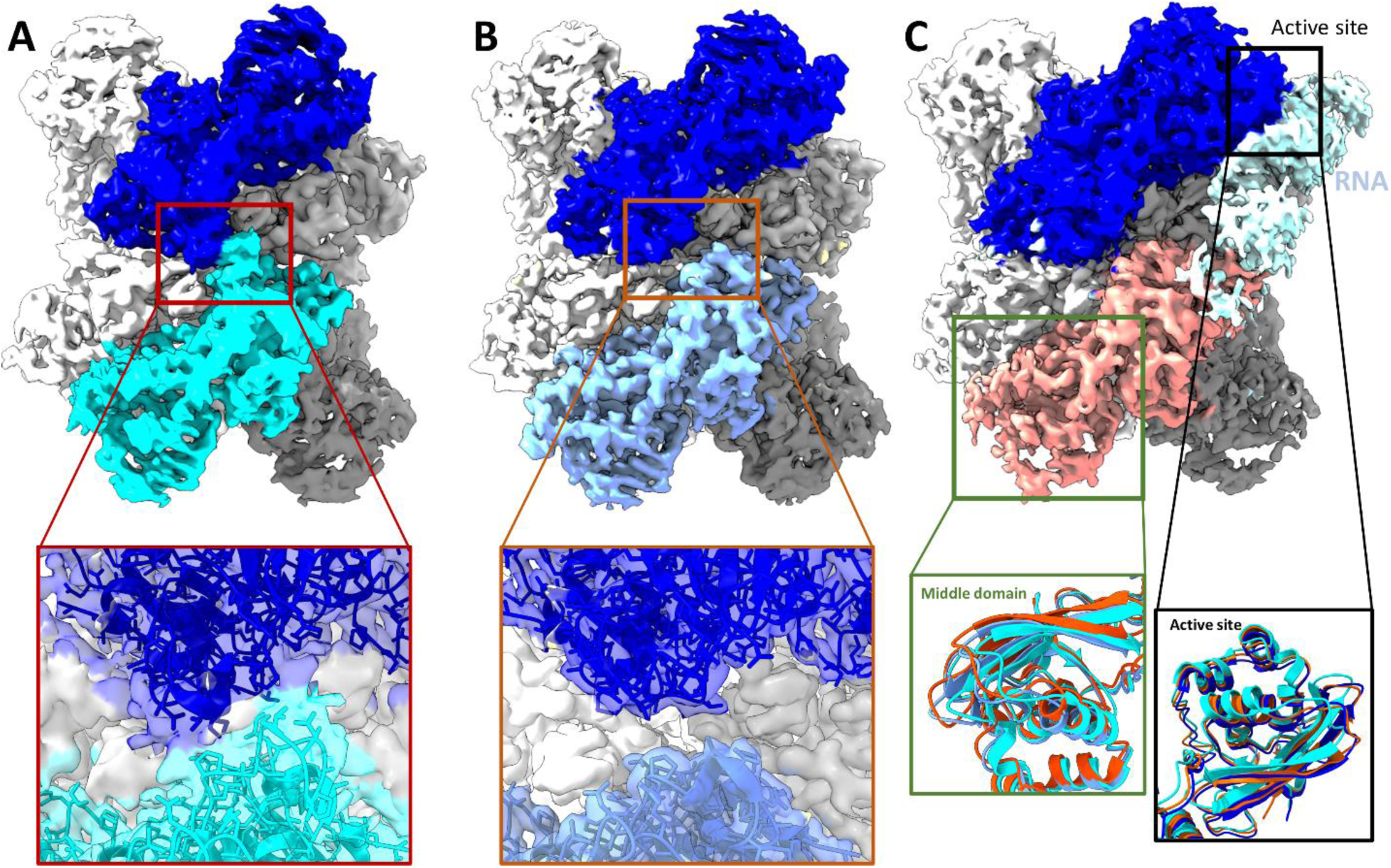
Cryo-EM maps of NendoU in different conditions. A) Cryo-EM maps of NendoU in pH 7.5 (PDB 7RB0) in closed state, with chains A and B collored in blue and cyan, respectively. Box shows detailed view of map and model from the switch region between these two chains. B) Cryo-EM maps of NendoU in pH 6.0 (PDB 7ME0) in open state, with chains A and B collored in blue and lightblue, respectively. Box shows detailed view of map and model from the switch region between these two chains. C) Cryo-EM maps of NendoU in complex with dsRNA (PDB 7TJ2) and in open state, with chains A and B collored in blue and salmon, respectively. dsRNA map is colored in lightblue. Black box shows detailed view of three models aligned in the active site region. Green box shows detailed views the three models middle domain as cartoons superposed in adjacent NendoU. 7RB0, 7ME0 and 7TJ2 are colored in dark blue, cyan and salmon, respectively. Maps are showed with contrast of 0.455.

In the NendoU dsRNA complex solved by Frazier *et al*. (*63*), the positively charged switch region can be found in the open form and seen to form key interactions with the nucleic acid, indicating that this region is involved in positioning the RNA into the active site in a form of the enzyme that resembles the open state (Fig. 2D). In fact, this open-closed form model seems to agree with our biochemical data showing that the enzyme is far more active in acidic pH than in basic, where the switch seems to be found in the closed conformation. In the dsRNA complex, we can also see that the MD active site region of the adjacent auxiliary NSP15 chain is twisted relative to the open and closed forms of NendoU (Fig 3C), while the occupied active site from the productive (bound to RNA in the active site) chain less altered among the states (Fig. 3C). In the movie generated by 3DVA of samples in HEPES pH 7.5 (Supplementary movie #1 and #2) and PBS pH 6.0 (Supplementary movie #3 and #4), we can see the wobbling movement of these chains relative to each other and highlight the conformational changes of the switch region, similar to the movements observed by Pilon *et al* (*11*). However, while they proposed the wobbling movement was the induced fit of enzyme processing mechanism, our data indicates that this movement is the shift between open-closed states of the enzyme, which regulate the enzyme active-inactive states, while the induced fit will occur with the conformational shifts of MD and switch regions within the constraints of the open state.

This might also explain why the structure of Frazier *et al*. (*63*) contains only one strand of nucleic acid at the same time, even that all other active sites are unoccupied during the binding, again agreeing with our biochemical characterization. This flexible mode of action involving two adjacent NSP15 chains interconnected by the switch region might explain not only why the cooperative index of NendoU is two (and not one, three or six), but also how and why pH affects the enzyme activity so drastically (Table S1). That model also diverges from recently proposed models in which bottom and top trimers would act independently in the substrate recognition and processing.

### NendoU polar surface allows stacking in the shape of filaments

A general observation we made was that particles from samples in basic pH’s, such as HEPES pH 7.5 (PDB 7RB0), tend to be individual particles (Fig. S9), while particles from samples obtained in more acid pH’s, such as BIS-Tris pH 6.0 (PDB 7RB2) and PBS pH 6.0 (PDB 7ME0), tend to be disposed side by side forming filaments (Fig. S10A), which is a common mechanism of allosteric control of enzyme activity (*65*). By using a large box size (512 × 512 px) for the sample in BIS-Tris pH 6.0, we were able to even classify and generate a low-resolution initial model of this dodecamer particle stack (Fig. S10B-C), which had also been observed in native mass spectrometry (Fig. S4). Using helical processing tools, we have been able to generate 2D classes of hand selected filaments, and data consistently indicate they are composed of C1 symmetrically paired units of the hexamer (Fig. 4A). The positioning of particles also indicate that hexamers are paired by the switch regions, but resolution of cryo-EM obtained model was not sufficient for describing these interactions (Fig. S10C). However, the most drastic similar effect was observed when we added the DNA oligoDT_30_ to the samples in BIS-Tris pH 6.0 and in PBS pH 6.0, where we saw the immediate formation of 2D crystals of NendoU in the frozen grid (Fig. S11). It seems that somehow the DNA in the sample causes NendoU particles to stack in this crystal pattern, which might be some mechanism of regulation.

**Fig 4.**
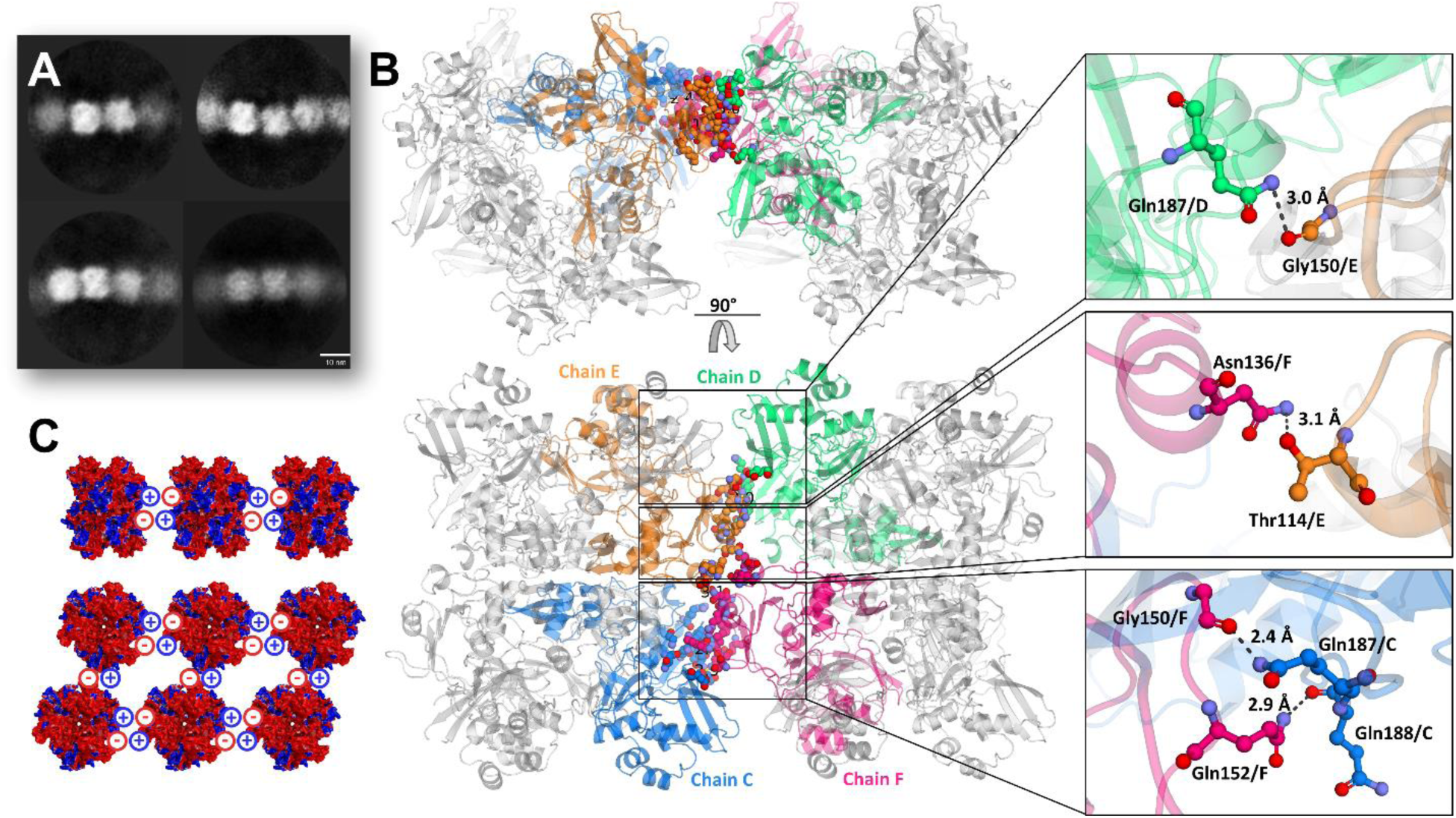
Overview of supramolecular organization of NendoU. A) 2D classification of selected filaments of NendoU observed from samples collected in BIS-Tris pH 6.0. B) Crystal structure of NendoU in presence of oligoDT reveals high resolution details of contacts between two NendoU hexamers switch region (PDB 7KF4). Protein is depictated as a cartoon, with chainds C, D, E and F colored in blue, green, orange and pink, respectively. Contacting residues are showed as colored sticks. C) Stacking model based on surface charge of NendoU. Structure of NendoU is colored according to its electrostatic potential projected on surface charge (-0.5 to 0.5 f kJ/mol/e in red-white-blue color model).

To further understand this stacking phenomenon, we tried to use this condition to obtain a crystal structure of NendoU. Typical crystals of NendoU are usually obtained in a hexagonal space group, with a dihedral asymmetric unit that can be symmetry expanded to generate the full hexamer. In these, the crystal packing resembles a chess board disposition of biological units, as an example of PDB’s 7KEG and 7KEH, with solvent content of about 74% (Fig. S12). However, when NendoU^mon^ was incubated with oligoDT, we observed that generated crystals were obtained in an orthorombic space group with 55% solvent content and biological units packed side by side as observed in the cryo-EM oligoDT samples, as exemplified here in PDB 7KF4 (Fig. S12). Furthermore, the same crystal pattern was also observed for NendoU^mon^ and NendoU^hex^ when incubated with other DNAs sequences such as primers (data not shown), suggesting that this packing is somehow induced by the presence of DNA. If we compare the disposition of adjacent hexamer units in the crystal structure (Fig. 4B) with the low-resolution processing of cryo-EM filaments (Fig. 4A and S10), one can infer that they follow the same pattern, with hexamer positioned side by side in a C1 disposition.

The high-resolution of the crystal structure (PDB 7KF4) allowed us to depict the contacts involved in the filament pattern, which involve the open form of the switch region of two adjacent hexamers, including the hydrogen bounds between Gln187 and Gly150, Asn136 and Thr114, Gly150 and Gnl187 and Gln152 and Gln188 (Fig. 4B). Furthermore, polar surface structural analysis showed that this pattern allows the perfect battery-like stacking of one hexamer negatively charged S cavity into the adjacent hexamer positively charged switch region (Fig. 2 and 4B), indicating that the biological function of these two regions is related to the formation of large order supramolecular agglomerates. To investigate the biological role of this still undescribed behavior, we generate the mutant T114L, which showed much higher activity in high concentrations of RNA substrate than WT (Fig. 1F), suggesting that this packing might be important for downregulating enzymatic activity in certain conditions. Yet, given the complexity of NendoU biochemical profile, we haven’t been able to fully understand the role of the switch region in enzyme regulation to date. More *in-situ* studies are required to confirm the existence of these molecular entities inside infect cells. Important to state that despite our efforts, we have not been able to identify any electron density suggesting a direct interaction between tested amino acids and NendoU.

### Crystallographic fragment screening of SARS-CoV-2 of NendoU

To identify starting points for new therapeutics, during the initial months of COVID-19 the XCHEM team and international collaborators have performed large crystallographic fragment screens against multiple SARS-CoV-2 proteins, including the viral Main protease, the Nsp3 macrodomain and the helicase Nsp13 (*66–68*), which had led the rapid development of new potential antivirals (*69*). Here we report the results obtained during the fragment screening campaign against NSP15 NendoU.

For the FBDD campaign, the chosen crystal system was obtained in a citrate condition and hexagonal space group *P*6_3_, containing two NSP15 monomers in the asymmetric unit disposed in a dihedral symmetry, which can be symmetrically expanded to reveal the full hexameric form of NendoU inside the crystal lattice. Three libraries of compounds were used for soaking, including the XCHEM Poised Library (*43*), the OPEN-EU DRIVE fragment library, and a set of 52 nucleoside analogs (Fig. 5). Over 1,200 crystals were soaked with different fragment compounds or nucleoside analogs, resulting in 997 datasets collected and processed, with an average resolution of 2.2 Å (Fig. 5B). Despite the high concentration of ligands used for soaking (100 mM), as well as the large amount and high quality of crystallographic data, only 26 crystal complexes of fragments or nucleosides (2.5%) with NendoU were obtained, highlighting the difficulties of identifying fragments for this crystal system, likely related to the large solvent content in the crystal unit cell (74%). From the 26 ligand complexes elucidated, we found that 13 ligands were found bounded to the two chains, while other 13 were only identified in one of the chains, which can be explained by the distinct dynamic between top and bottom trimers, related to the mechanism of action of NendoU (*11*). The average molecular weight (MW) of fragments identified was 183 ± 43 Da. The summarized information about PDB accession codes for each complex, resolution, refinement and processing statistics, ligand structure and chain location are available in Supporting table 1.

**Fig. 5.**
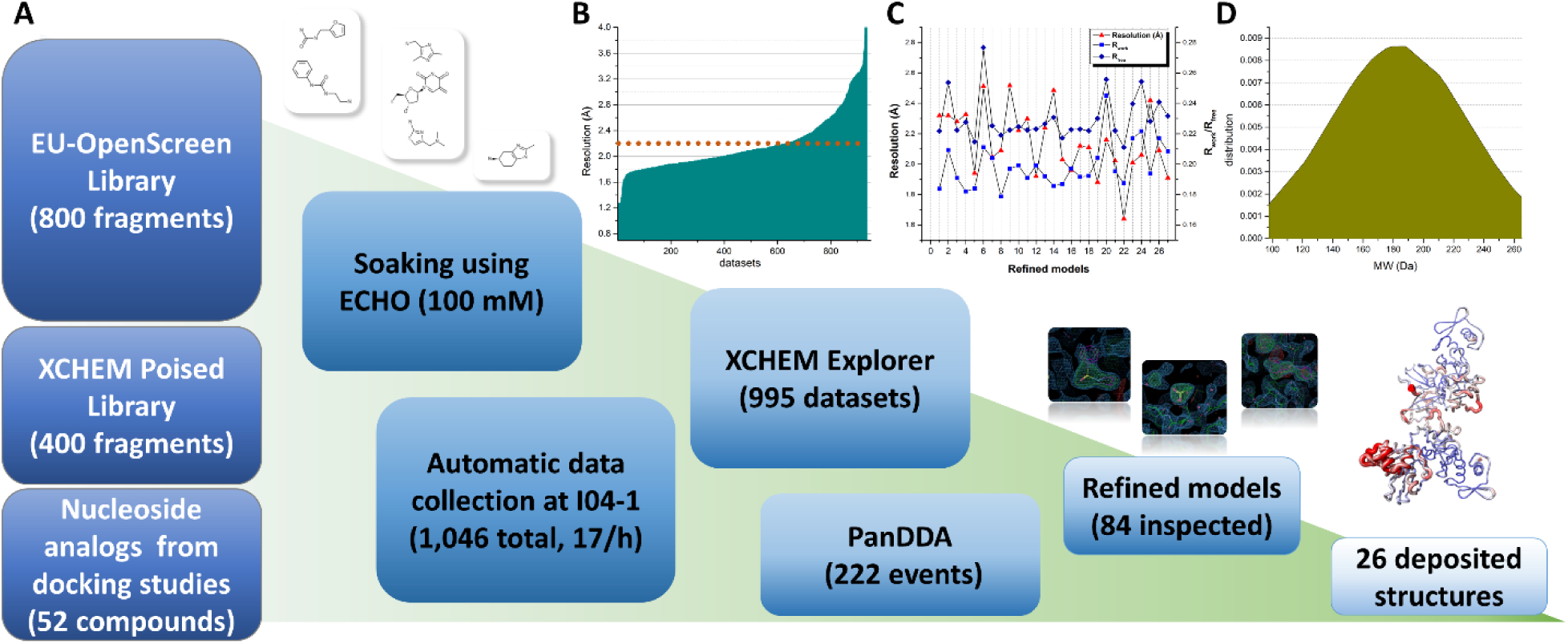
Overall schematic showing steps and data processing of the fragment screening campaing against NendoU. A) Schematic summarizing steps from library screening to refinement and deposition of models. Panels show event maps of selected fragments (1.0 sigma) and the overlap of all structures colored according to b-factors. B) Graph showing obtained resolution per dataset collected and processed. Doted line marks average resolution of all 997 usable datasets (2.2 Å). C) graph showing resolution and refinement statistics of final models deposited. Resolution is colored in red triangles, R_work_ is colored in blue squares and R_free_ is colored in dark blue diamonds. D) Normal distribution graph of molecular weitght from identified fragments.

To simplify our analyses, we superposed chains A and B of each complex onto chain A of a hexamer structure (PDBid 7KF4), so that we can map all the identified druggable cavities in the context of the hexameric complex (Fig. 6A). Also, we used the recent structure of NendoU in complex with double-strand RNA (PDBid 7TJ2) to inspect any of these sites that are in the path of the RNA binding surface (*63*). We found fourteen distinct binding sites spread across the NendoU surface, one at the ND, nine at the MD, and four at the CD. In the hexamer context, these sites are distributed across all the external and internal surfaces of the complex (Fig. 6A). Among those, we identified four sites where two or more ligands have formed clusters, which opens the possibility of fragment expansion.

**Fig 6.**
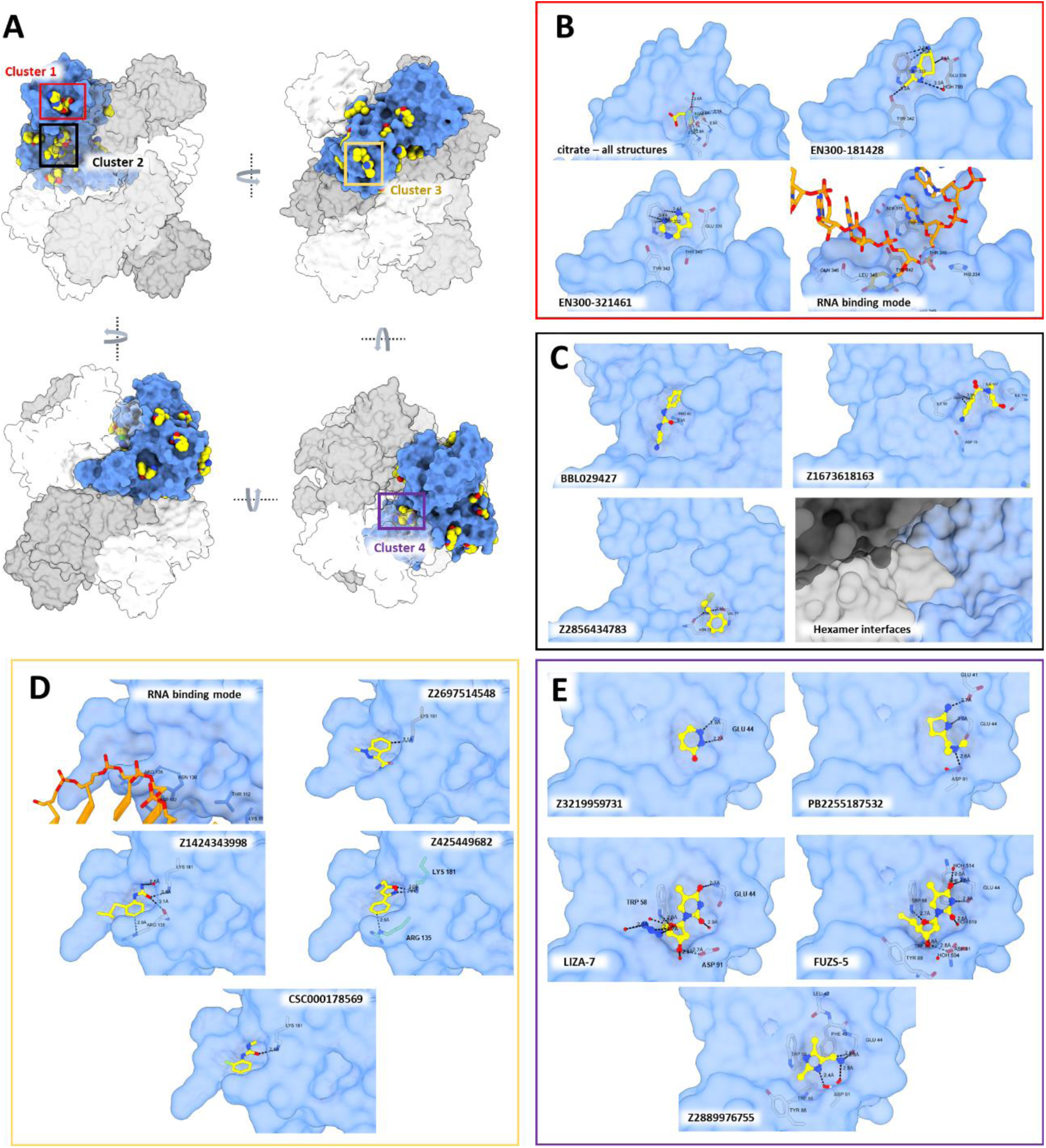
Overall visualization of fragments bound to NendoU. A) Overall view of all NendoU sites bound to fragments in multiple orientations. All chains containing fragments were supperposed into one, collored as blue surface, while adjacent chains of biological unit are collored in shades of grey. Colored squares highlight cluster of fragments identified. B) Detailed view of cluster 1 (active site) showing citrate, multiple fragment contacts and RNA binding mode. C) Detailed view of cluster 2 showing multiple fragment contacts and oligomerization interface. D) Detailed view of cluster 3, showing multiple fragment contacts and RNA binding mode. E) Detailed view of cluster 4, showing multiple fragment contacts with NendoU. In all, NendoU surface is collored in blue, fragments are depicted as sticks or spheres with yellow carbons, while interacting NendoU residues are depicted as grey carbon sticks. RNA carbons are collored in orange, and coordinates were obtained from PDB 7TJ2.

Unfortunately, the NendoU active site, which would be a primary target for the development of a competitive inhibitor, was occupied by a citrate molecule that originated from the crystallization condition in all refined datasets (Fig. 6B), which resulted in a small number of binders to this site. The only identified fragments in this site were two aromatic fragments, EN300-181428 and EN300-321461, forming a π-π stacking interaction with Trp332 (Fig. 6B). This is consistent with the mode of action of NendoU, which prefers an aromatic purine at the +2 position together with the intrinsic requirement of a uridine in the cleavage position.

The second cluster (Cluster 2) mapped is in a conserved region of the MD, which is a critical region of the oligomerization interface (Fig. 6C). Here we identified a large cavity in which fragments BBL029427, Z1673618163 and Z2856434783 are forming hydrogen bonds with multiple MD residues, including Pro93, Ser97, Asn73 and Val77 (Fig. 6C). These three fragments are spread across the cavity, permitting a good exploration of its chemical space and potential for fragment linking or merging.

Another possible allosteric site for the development of new inhibitors is the one identified in the third cluster (Cluster 3), which is also a conserved region located at the bottom of the MD, where we found a cavity containing fragments Z2697514548, Z1424343998, Z425449682 and CSC000178569, forming different contacts with residues Arg135 and Lys191 (Fig. 6D). The structure of NendoU in complex with dsRNA reveals that the positive charges of Arg135 and Lys191 sidechains are key for positioning long nucleic acids into the active sites of NendoU (*63*).

The most curious site identified in our experiments was cluster 4, which is a conserved site located in the interfaces of ND and MD to the NSP15 domain of the neighboring monomer, in a cavity located inside the barrel shape of NendoU (Fig. 6E). Beside the aromatic fragments Z3219959731, PB2255187532 and Z2889976755, we also identified two nucleoside analogs, LIZA-7 and FUZS-5, tightly bound to this allosteric site (Fig. 6E). The two thymidine analogs are 5’-azido-5’-deoxy-thymidine (LIZA-7) and 5’-deoxy-5’-thiothymidine (FUZS-5), with their deoxyribose unit showing two productive hydrogen bonds with Trp58 and Asp91, as well as a π-π stacking between the nucleoside base and Trp58. To further investigate the role of this allosteric site, we have examined the effect of FUZS-5 on the stability and enzymatic activity of NSP15, with a thermal stability assay, and a fluorescence-based enzyme activity assay, respectively. Notably, FUZS-5 has increased the thermal stability of NSP15 by 0.4 °C, suggesting a stabilizing role for the compound itself, as well as other potential binders of this allosteric site. Furthermore, the enzyme activity assay revealed FUZS-5 to increase the reaction rate, in a closely linear, concentration-dependent fashion (Fig. S13). Together with the X-ray structure, these results suggest the positive allosteric modulation of NendoU by FUZS-5, pointing at a previously unknown, positive feedback mechanism for the activity of NSP15. The close structural analogy of FUZS-5 to the natural nucleoside thymidine (as well as uridine) suggests that this positive modulatory mechanism might be triggered under physiological conditions as well, by the endogenous nucleosides. More studies are being conducted to clarify this mechanism.

Besides the four identified clusters, we were also able to map several different druggable cavities spreads across the hexamer structure, including fragments EN300-1605072, Z2443429438, Z425449682, Z59181945, Z239136710, Z1530301542, Z18197050, EN300-100112, Z319891284, Z68299550, Z56900771 and Z31504642, which are all depicted in Fig. 7. In summary, our data allowed us to map several new sites for the development of allosteric inhibitors that had never been described for NendoU, including multiples sites that might be used block oligomerization or RNA recognition (Fig. 6A). The undergoing expansion of selected fragments and characterization will be critical for validating the therapeutic potential of these sites, opening opportunities for the development of new allosteric inhibitors of NendoU.

**Fig. 7.**
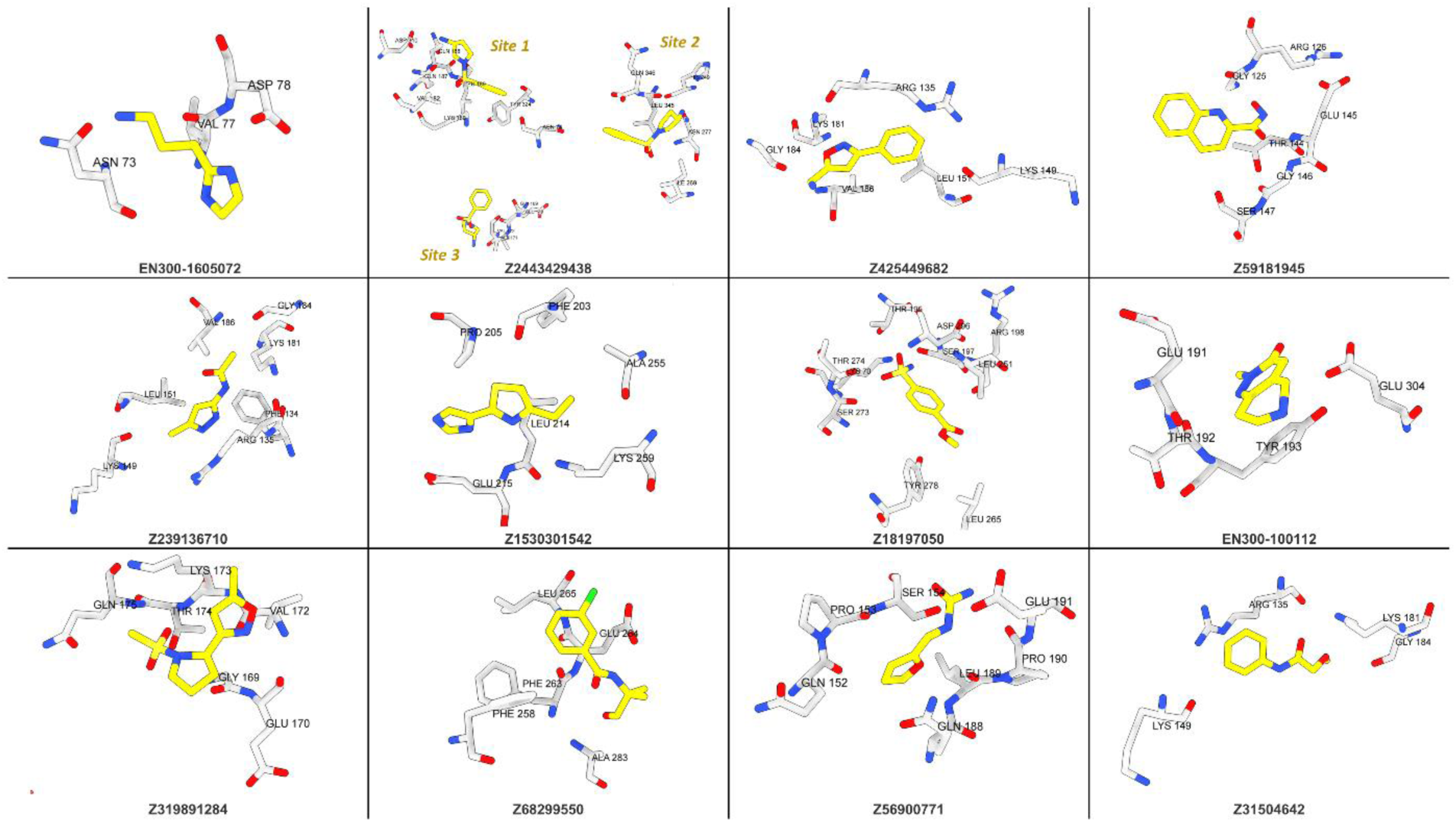
Other fragments identified during fragment screening spread accros multiple putative allosteric sites. Fragments are depicted as sticks with yellow carbons, while interacting NendoU residues are depicted as grey carbon sticks.

### Fragments caused the disruption of crystal structure from NendoU active site

During our analysis of fragment screening data, we identified a set of datasets that had a completely disrupted chain A active site, causing an extreme increase in the B-factor of this region (Fig. 8A). A detailed inspection of these datasets showed no significant twinning or other crystal pathology that would explain such poor electron density. Yet, despite our efforts we were not able to obtain a refined map of this region, or identify fragments bound to that region. Still, given all the previous observations related to mobility of one individual active site and the lack of fragments bound to the active, it is likely that these fragments are somehow interacting with the active site region and causing the disruption of the processive or auxiliary chains only. These are fragments Z1741794237, Z1713595338, Z85934875, Z1437171658, Z1636723439, Z2856434903, Z1262327505, Z1269638430, Z927400026 and POB0120, and their structures are depicted in Fig. 8B. The datasets of these fragments were not deposited in PDB, but can be found in Supplementary datasets file. More studies are required to understand the biological impact of these fragments in NendoU activity and define their mode of interaction.

**Fig. 8.**
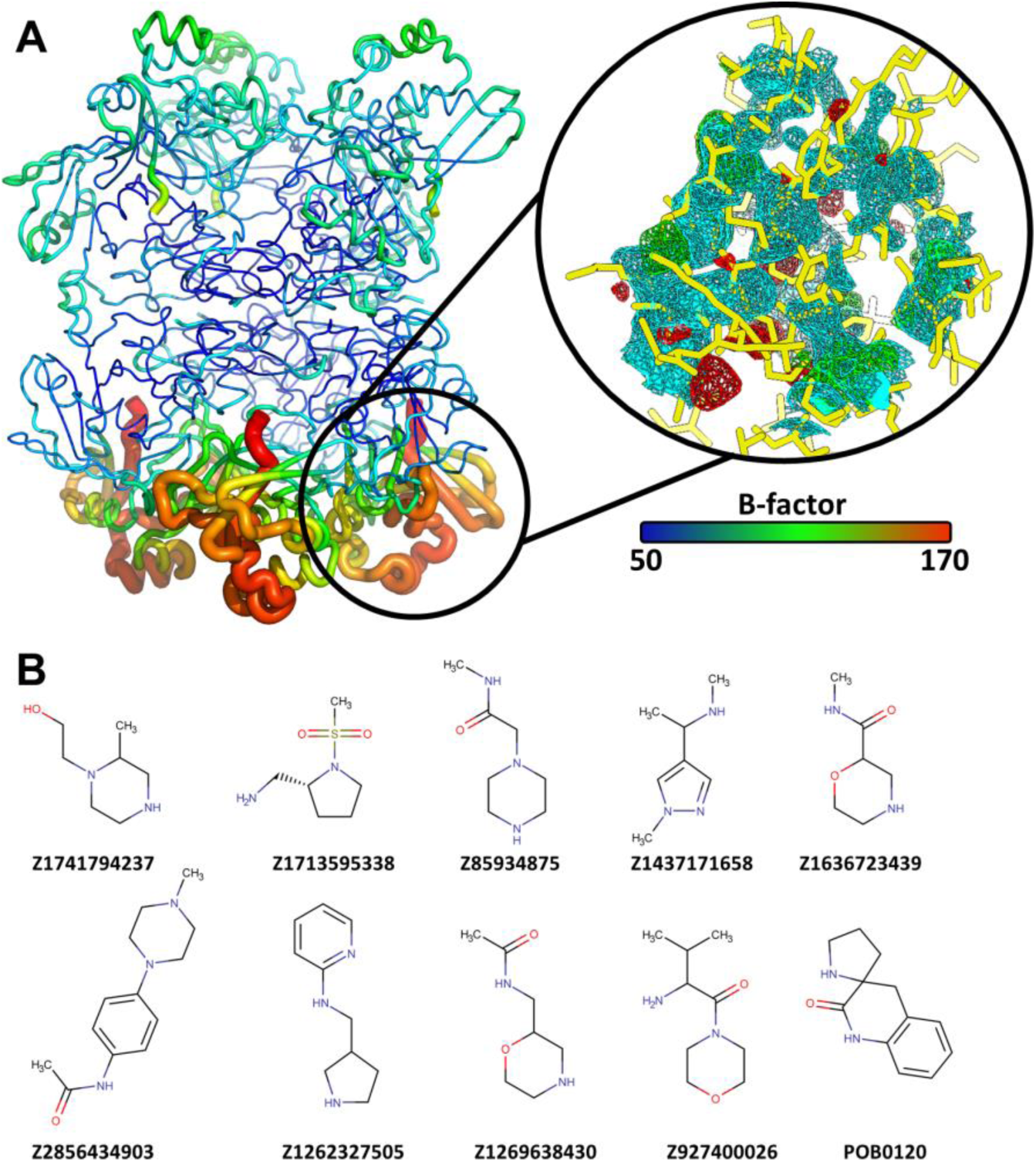
Fragments that caused disruption of active site of one chain. A) View from hexamer colored according to its B-factor (50 to 170 Å^2^), highlighting the oberserved effect on chain A. The highlighted region shows chain A activite site region with one example (dataset #134) of the electron density observed for this region. 2Fo-Fc map is colored in cyan, while Fo-Fc maps are colored in green (positive values) or red (negative values). B) Chemical structure of all fragments identified that caused disruption of chain A active site.

## AVAILABILITY

Any other data can be supplied upon reasonable request.

## ACCESSION NUMBERS

Crystallographic coordinates and structure factors for all structures have been deposited in the PDB with the following accessing codes 5S6X, 5S6Y, 5S6Z, 5S70, 5S71, 5S72, 5SA4, 5SA5, 5SA6, 5SA7, 5SA8, 5SA9, 5SAA, 5SAB, 5SAC, 5SAD, 5SAE, 5SAF, 5SAG, 5SAH, 5SAI, 5SBF, 7N7R, 7N7U, 7N7W, 7N7Y, 7N83, 7KEG, 7KF4, 7KEH, 7RB0, 7RB2 and 7ME0. EMD data is available with the following accessing codes EMD-24391, EMD-24392, EMD-23786.

## SUPPLEMENTARY DATA

Supplementary Data are available at NAR online.

## ACKNOWLEDGEMENT

Authors acknowledge Diamond Light Source for access and support of the cryo-EM facilities at the UK national electron Bio-Imaging Centre (eBIC) through proposal BI27083, funded by the Wellcome Trust, MRC and BBSRC, and for access to the fragment screening facility XChem, for usage of DSi-Poised library and for beamtime on beamline I04-1 under proposal LB27023. This research used facilities of the SIRIUS, part of the Brazilian Center for Research in Energy and Materials (CNPEM), a private non-profit organization under the supervision of the Brazilian Ministry for Science, Technology, and Innovations (MCTI). The MANACA beamline staff is acknowledged for the assistance during the experiments of proposal 20200014. We acknowledge MAX IV Laboratory for time on Beamline BioMAX under Proposal MX20190328. Research conducted at MAX IV, a Swedish national user facility, is supported by the Swedish Research council under contract 2018-07152, the Swedish Governmental Agency for Innovation Systems under contract 2018-04969, and Formas under contract 2019-02496. Authors acknowledges EU-OPENSCREEN ERIC for providing its fragment library for the presented scientific work. EU-OPENSCREEN ERIC has received funding from European Union’s Horizon 2020 research and innovation program under grant agreement No 823893 (EUOPENSCREEN-DRIVE).

## References

1. P. Zhou, X. Lou Yang, X. G. Wang, B. Hu, L. Zhang, W. Zhang, H. R. Si, Y. Zhu, B. Li, C. L. Huang, H. D. Chen, J. Chen, Y. Luo, H. Guo, R. Di Jiang, M. Q. Liu, Y. Chen, X. R. Shen, X. Wang, X. S. Zheng, K. Zhao, Q. J. Chen, F. Deng, L. L. Liu, B. Yan, F. X. Zhan, Y. Y. Wang, G. F. Xiao, Z. L. Shi, A pneumonia outbreak associated with a new coronavirus of probable bat origin. Nature. 579, 270–273 (2020).

2. N. Zhu, D. Zhang, W. Wang, X. Li, B. Yang, J. Song, X. Zhao, B. Huang, W. Shi, R. Lu, P. Niu, F. Zhan, X. Ma, D. Wang, W. Xu, G. Wu, G. F. Gao, W. Tan, A novel coronavirus from patients with pneumonia in China, 2019. New England Journal of Medicine. 382, 727–733 (2020).

3. H. Li, Y. Zhou, M. Zhang, H. Wang, Q. Zhao, J. Liu, Updated approaches against SARS-CoV-2. Antimicrob Agents Chemother. 64 (2020), pp. 1–7.

4. R. Lu, X. Zhao, J. Li, P. Niu, B. Yang, H. Wu, W. Wang, H. Song, B. Huang, N. Zhu, Y. Bi, X. Ma, F. Zhan, L. Wang, T. Hu, H. Zhou, Z. Hu, W. Zhou, L. Zhao, J. Chen, Y. Meng, J. Wang, Y. Lin, J. Yuan, Z. Xie, J. Ma, W. J. Liu, D. Wang, W. Xu, E. C. Holmes, G. F. Gao, G. Wu, W. Chen, W. Shi, W. Tan, Genomic characterisation and epidemiology of 2019 novel coronavirus: implications for virus origins and receptor binding. The Lancet (2020), doi:10.1016/S0140-6736(20)30251-8.

5. H. Li, Z. Liu, J. Ge, Scientific research progress of COVID-19/SARS-CoV-2 in the first five months. J Cell Mol Med. 24, 6558 (2020).

6. P. R. Bhatt, A. Scaiola, G. Loughran, M. Leibundgut, A. Kratzel, R. Meurs, R. Dreos, K. M. O’Connor, A. McMillan, J. W. Bode, V. Thiel, D. Gatfield, J. F. Atkins, N. Ban, Structural basis of ribosomal frameshifting during translation of the SARS-CoV-2 RNA genome. Science (1979). 372, 1306–1313 (2021).

7. S. L, I. I, F. F, C. A, C. B, D. E, C. B, SARS-CoV ORF1b-encoded nonstructural proteins 12-16: replicative enzymes as antiviral targets. Antiviral Res. 101, 122–130 (2014).

8. Y. Kim, R. Jedrzejczak, N. I. Maltseva, M. Wilamowski, M. Endres, A. Godzik, K. Michalska, A. Joachimiak, Crystal structure of Nsp15 endoribonuclease NendoU from SARS-CoV-2. Protein Science. 29, 1596–1605 (2020).

9. E. J. Snijder, P. J. Bredenbeek, J. C. Dobbe, V. Thiel, J. Ziebuhr, L. L. M. Poon, Y. Guan, M. Rozanov, W. J. M. Spaan, A. E. Gorbalenya, Unique and Conserved Features of Genome and Proteome of SARS-coronavirus, an Early Split-off From the Coronavirus Group 2 Lineage. J Mol Biol. 331, 991– 1004 (2003).

10. K. Bhardwaj, S. Palaninathan, J. M. O. Alcantara, L. L. Yi, L. Guarino, J. C. Sacchettini, C. C. Kao, Structural and functional analyses of the severe acute respiratory syndrome coronavirus endoribonuclease Nsp15. Journal of Biological Chemistry. 283, 3655–3664 (2008).

11. M. C. Pillon, M. N. Frazier, L. B. Dillard, J. G. Williams, S. Kocaman, J. M. Krahn, L. Perera, C. K. Hayne, J. Gordon, Z. D. Stewart, M. Sobhany, L. J. Deterding, A. L. Hsu, V. P. Dandey, M. J. Borgnia, R. E. Stanley, Cryo-EM structures of the SARS-CoV-2 endoribonuclease Nsp15 reveal insight into nuclease specificity and dynamics. Nat Commun. 12, 1–12 (2021).

12. L. A. Guarino, K. Bhardwaj, W. Dong, J. Sun, A. Holzenburg, C. Kao, Mutational analysis of the SARS virus Nsp15 endoribonuclease: Identification of residues affecting hexamer formation. J Mol Biol. 353, 1106–1117 (2005).

13. S. T. Shi, J. J. Schiller, A. Kanjanahaluethai, S. C. Baker, J.-W. Oh, M. M. C. Lai, Colocalization and Membrane Association of Murine Hepatitis Virus Gene 1 Products and De Novo-Synthesized Viral RNA in Infected Cells. J Virol. 73, 5957–5969 (1999).

14. J. Athmer, A. R. Fehr, M. Grunewald, E. C. Smith, M. R. Denison, S. Perlman, In Situ Tagged nsp15 Reveals Interactions with Coronavirus Replication/Transcription Complex-Associated Proteins. mBio. 8 (2017), doi:10.1128/MBIO.02320-16.

15. L. Yan, Y. Zhang, J. Ge, L. Zheng, Y. Gao, T. Wang, Z. Jia, H. Wang, Y. Huang, M. Li, Q. Wang, Z. Rao, Z. Lou, Architecture of a SARS-CoV-2 mini replication and transcription complex. Nature Communications 2020 11:1. 11, 1–6 (2020).

16. X. Deng, M. Hackbart, R. C. Mettelman, A. O’Brien, A. M. Mielech, G. Yi, C. C. Kao, S. C. Baker, Coronavirus nonstructural protein 15 mediates evasion of dsRNA sensors and limits apoptosis in macrophages. Proceedings of the National Academy of Sciences. 114, E4251–E4260 (2017).

17. X. Deng, S. C. Baker, An “Old” protein with a new story: Coronavirus endoribonuclease is important for evading host antiviral defenses. Virology. 517, 157–163 (2018).

18. C.-K. Yuen, J.-Y. Lam, W.-M. Wong, L.-F. Mak, X. Wang, H. Chu, J.-P. Cai, D.-Y. Jin, K. K.-W. To, J. F.-Chan, K.-Y. Yuen, K.-H. Kok, SARS-CoV-2 nsp13, nsp14, nsp15 and orf6 function as potent interferon antagonists. https://doi.org/10.1080/22221751.2020.1780953. 9, 1418–1428 (2020).

19. E. Kindler, C. Gil-Cruz, J. Spanier, Y. Li, J. Wilhelm, H. H. Rabouw, R. Züst, M. Hwang, P. V’kovski, H. Stalder, S. Marti, M. Habjan, L. Cervantes-Barragan, R. Elliot, N. Karl, C. Gaughan, F. J. M. van Kuppeveld, R. H. Silverman, M. Keller, B. Ludewig, C. C. Bergmann, J. Ziebuhr, S. R. Weiss, U. Kalinke, V. Thiel, Early endonuclease-mediated evasion of RNA sensing ensures efficient coronavirus replication. PLoS Pathog. 13, e1006195 (2017).

20. D. E. Gordon, G. M. Jang, M. Bouhaddou, J. Xu, K. Obernier, K. M. White, M. J. O’Meara, V. V. Rezelj, J. Z. Guo, D. L. Swaney, T. A. Tummino, R. Hüttenhain, R. M. Kaake, A. L. Richards, B. Tutuncuoglu, H. Foussard, J. Batra, K. Haas, M. Modak, M. Kim, P. Haas, B. J. Polacco, H. Braberg, J. M. Fabius, M. Eckhardt, M. Soucheray, M. J. Bennett, M. Cakir, M. J. McGregor, Q. Li, B. Meyer, F. Roesch, T. Vallet, A. Mac Kain, L. Miorin, E. Moreno, Z. Z. C. Naing, Y. Zhou, S. Peng, Y. Shi, Z. Zhang, W. Shen, I. T. Kirby, J. E. Melnyk, J. S. Chorba, K. Lou, S. A. Dai, I. Barrio-Hernandez, D. Memon, C. Hernandez-Armenta, J. Lyu, C. J. P. Mathy, T. Perica, K. B. Pilla, S. J. Ganesan, D. J. Saltzberg, R. Rakesh, X. Liu, S. B. Rosenthal, L. Calviello, S. Venkataramanan, J. Liboy-Lugo, Y. Lin, P. Huang, Y. F. Liu, S. A. Wankowicz, M. Bohn, M. Safari, F. S. Ugur, C. Koh, N. S. Savar, Q. D. Tran, D. Shengjuler, S. J. Fletcher, M. C. O’Neal, Y. Cai, J. C. J. Chang, D. J. Broadhurst, S. Klippsten, P. P. Sharp, N. A. Wenzell, D. Kuzuoglu-Ozturk, H. Y. Wang, R. Trenker, J. M. Young, D. A. Cavero, J. Hiatt, T. L. Roth, U. Rathore, A. Subramanian, J. Noack, M. Hubert, R. M. Stroud, A. D. Frankel, O. S. Rosenberg, K. A. Verba, D. A. Agard, M. Ott, M. Emerman, N. Jura, M. von Zastrow, E. Verdin, A. Ashworth, O. Schwartz, C. d’Enfert, S. Mukherjee, M. Jacobson, H. S. Malik, D. G. Fujimori, T. Ideker, C. S. Craik, S. N. Floor, J. S. Fraser, J. D. Gross, A. Sali, B. L. Roth, D. Ruggero, J. Taunton, T. Kortemme, P. Beltrao, M. Vignuzzi, A. García-Sastre, K. M. Shokat, B. K. Shoichet, N. J. Krogan, A SARS-CoV-2 protein interaction map reveals targets for drug repurposing. Nature. 583, 459–468 (2020).

21. M. Hackbart, X. Deng, S. C. Baker, Coronavirus endoribonuclease targets viral polyuridine sequences to evade activating host sensors. Proceedings of the National Academy of Sciences. 117, 8094–8103 (2020).

22. C. Aslanidis, P. J. de Jong, Ligation-independent cloning of PCR products (LIC-PCR). Nucleic Acids Res. 18, 6069–6074 (1990).

23. M. R. Wilkins, E. Gasteiger, A. Bairoch, J. C. Sanchez, K. L. Williams, R. D. Appel, D. F. Hochstrasser, Protein identification and analysis tools in the ExPASy server. Methods Mol Biol. 112, 531–552 (1999).

24. J. Ortiz-Alcantara, K. Bhardwaj, S. Palaninathan, M. Frieman, R. S. Baric, C. C. Kao, Small molecule inhibitors of the SARS-CoV Nsp15 endoribonuclease. Virus Adaptation and Treatment. 2, 125–133 (2010).

25. S. Q. Zheng, E. Palovcak, J. P. Armache, K. A. Verba, Y. Cheng, D. A. Agard, MotionCor2: Anisotropic correction of beam-induced motion for improved cryo-electron microscopy. Nat Methods. 14 (2017), pp. 331–332.

26. A. Punjani, J. L. Rubinstein, D. J. Fleet, M. A. Brubaker, CryoSPARC: Algorithms for rapid unsupervised cryo-EM structure determination. Nat Methods. 14, 290–296 (2017).

27. P. v. Afonine, B. K. Poon, R. J. Read, O. v. Sobolev, T. C. Terwilliger, A. Urzhumtsev, P. D. Adams, Real-space refinement in PHENIX for cryo-EM and crystallography. Acta Crystallogr D Struct Biol. 74, 531–544 (2018).

28. P. Ef, G. Td, H. Cc, M. Ec, C. Gs, C. Ti, M. Jh, F. Te, UCSF ChimeraX: Structure visualization for researchers, educators, and developers. Protein Sci. 30, 70–82 (2021).

29. A. Punjani, D. J. Fleet, 3D variability analysis: Resolving continuous flexibility and discrete heterogeneity from single particle cryo-EM. J Struct Biol. 213, 107702 (2021).

30. J. Zivanov, T. Nakane, B. O. Forsberg, D. Kimanius, W. J. H. Hagen, E. Lindahl, S. H. W. Scheres, New tools for automated high-resolution cryo-EM structure determination in RELION-3. Elife. 7 (2018), doi:10.7554/ELIFE.42166.

31. Z. Sq, P. E A. Jp, V. Ka, C. Y A. Da, MotionCor2: anisotropic correction of beam-induced motion for improved cryo-electron microscopy. Nat Methods. 14, 331–332 (2017).

32. R. A, G. N, CTFFIND4: Fast and accurate defocus estimation from electron micrographs. J Struct Biol. 192, 216–221 (2015).

33. W. Kabsch, XDS. Acta Crystallogr D Biol Crystallogr. 66, 125–132 (2010).

34. C. Vonrhein, C. Flensburg, P. Keller, A. Sharff, O. Smart, W. Paciorek, T. Womack, G. Bricogne, Data processing and analysis with the autoPROC toolbox. Acta Crystallogr D Biol Crystallogr. 67, 293–302 (2011).

35. M. D. Winn, C. C. Ballard, K. D. Cowtan, E. J. Dodson, P. Emsley, P. R. Evans, R. M. Keegan, E. B. Krissinel, A. G. W. Leslie, A. McCoy, S. J. McNicholas, G. N. Murshudov, N. S. Pannu, E. A. Potterton, H. R. Powell, R. J. Read, A. Vagin, K. S. Wilson, Overview of the CCP4 suite and current developments. Acta Crystallographica Section D. 67, 235–242 (2011).

36. A. J. McCoy, R. W. Grosse-Kunstleve, P. D. Adams, M. D. Winn, L. C. Storoni, R. J. Read, Phaser crystallographic software. J Appl Crystallogr. 40, 658–674 (2007).

37. P. v Afonine, R. W. Grosse-Kunstleve, N. Echols, J. J. Headd, N. W. Moriarty, M. Mustyakimov, T. C. Terwilliger, A. Urzhumtsev, P. H. Zwart, P. D. Adams, Towards automated crystallographic structure refinement with phenix.refine. Acta Crystallographica Section D. 68, 352–367 (2012).

38. O. S. Smart, T. O. Womack, C. Flensburg, P. Keller, W. Paciorek, A. Sharff, C. Vonrhein, G. Bricogne, Exploiting structure similarity in refinement: Automated NCS and target-structure restraints in BUSTER. Acta Crystallogr D Biol Crystallogr. 68, 368–380 (2012).

39. P. Emsley, B. Lohkamp, W. G. Scott, K. Cowtan, Features and development of Coot. Acta Crystallogr D Biol Crystallogr. 66, 486–501 (2010).

40. V. B. Chen, W. B. Arendall 3rd, J. J. Headd, D. A. Keedy, R. M. Immormino, G. J. Kapral, L. W. Murray, J. S. Richardson, D. C. Richardson, MolProbity: all-atom structure validation for macromolecular crystallography. Acta Crystallogr D Biol Crystallogr. 66, 12–21 (2010).

41. E. Jurrus, D. Engel, K. Star, K. Monson, J. Brandi, L. E. Felberg, D. H. Brookes, L. Wilson, J. Chen, K. Liles, M. Chun, P. Li, D. W. Gohara, T. Dolinsky, R. Konecny, D. R. Koes, J. E. Nielsen, T. Head-Gordon, W. Geng, R. Krasny, G. W. Wei, M. J. Holst, J. A. McCammon, N. A. Baker, Improvements to the APBS biomolecular solvation software suite. Protein Science. 27, 112–128 (2018).

42. H. Ashkenazy, S. Abadi, E. Martz, O. Chay, I. Mayrose, T. Pupko, N. Ben-Tal, ConSurf 2016: an improved methodology to estimate and visualize evolutionary conservation in macromolecules. Nucleic Acids Res. 44, W344–50 (2016).

43. O. B. Cox, T. Krojer, P. Collins, O. Monteiro, R. Talon, A. Bradley, O. Fedorov, J. Amin, B. D. Marsden, J. Spencer, F. von Delft, P. E. Brennan, A poised fragment library enables rapid synthetic expansion yielding the first reported inhibitors of PHIP(2), an atypical bromodomain. Chem Sci. 7, 2322–2330 (2016).

44. P. M. Collins, J. T. Ng, R. Talon, K. Nekrosiute, T. Krojer, A. Douangamath, J. Brandao-Neto, N. Wright, N. M. Pearce, F. Von Delft, Gentle, fast and effective crystal soaking by acoustic dispensing. Acta Crystallogr D Struct Biol. 73, 246–255 (2017).

45. S. E. Thomas, P. Collins, R. H. James, V. Mendes, S. Charoensutthivarakul, C. Radoux, C. Abell, A. G. Coyne, R. A. Floto, F. von Delft, T. L. Blundell, Structure-guided fragment-based drug discovery at the synchrotron: screening binding sites and correlations with hotspot mapping. Philosophical Transactions of the Royal Society A: Mathematical, Physical and Engineering Sciences. 377, 20180422 (2019).

46. M. D. Winn, C. C. Ballard, K. D. Cowtan, E. J. Dodson, P. Emsley, P. R. Evans, R. M. Keegan, E. B. Krissinel, A. G. W. Leslie, A. McCoy, S. J. McNicholas, G. N. Murshudov, N. S. Pannu, E. A. Potterton, H. R. Powell, R. J. Read, A. Vagin, K. S. Wilson, Overview of the CCP4 suite and current developments. Acta Crystallographica Section D. 67, 235–242 (2011).

47. N. M. Pearce, T. Krojer, A. R. Bradley, P. Collins, R. P. Nowak, R. Talon, B. D. Marsden, S. Kelm, J. Shi, C. M. Deane, F. Von Delft, A multi-crystal method for extracting obscured crystallographic states from conventionally uninterpretable electron density. Nat Commun. 8 (2017), doi:10.1038/ncomms15123.

48. H. M. Ginn, Pre-clustering data sets using cluster4x improves the signal-to-noise ratio of high-throughput crystallography drug-screening analysis. urn:issn:2059-7983. 76, 1134–1144 (2020).

49. R. Winkler, ESIprot: a universal tool for charge state determination and molecular weight calculation of proteins from electrospray ionization mass spectrometry data. Rapid Communications in Mass Spectrometry. 24, 285–294 (2010).

50. N. Debreczeni, M. Bege, M. Herczeg, I. Bereczki, G. Batta, P. Herczegh, A. Borbás, Tightly linked morpholino-nucleoside chimeras: new, compact cationic oligonucleotide analogues. Org Biomol Chem. 19, 8711–8721 (2021).

51. I. Van Daele, H. Munier-Lehmann, M. Froeyen, J. Balzarini, S. Van Calenbergh, Rational design of 5′-thiourea-substituted α-thymidine analogues as thymidine monophosphate kinase inhibitors capable of inhibiting mycobacterial growth. J Med Chem. 50, 5281–5292 (2007).

52. S. H. Kawai, D. Wang, G. Just, Synthesis of the thymidine building blocks for a nonhydrolyzable DNA analogue. Can J Chem. 70, 1573–1580 (1992).

53. E. J. Reist, A. Benitez, L. Goodman, The Synthesis of Some 5’-Thiopentofuranosylpyrimidines1. Journal of Organic Chemistry. 29, 554–558 (2002).

54. K. Bhardwaj, J. Sun, A. Holzenburg, L. A. Guarino, C. C. Kao, RNA Recognition and Cleavage by the SARS Coronavirus Endoribonuclease. J Mol Biol. 361, 243–256 (2006).

55. F. H. Niesen, H. Berglund, M. Vedadi, The use of differential scanning fluorimetry to detect ligand interactions that promote protein stability. Nat Protoc. 2, 2212–2221 (2007).

56. K. Bhardwaj, J. Sun, A. Holzenburg, L. A. Guarino, C. C. Kao, RNA Recognition and Cleavage by the SARS Coronavirus Endoribonuclease. J Mol Biol. 361, 243–256 (2006).

57. L. A. Guarino, K. Bhardwaj, W. Dong, J. Sun, A. Holzenburg, C. Kao, Mutational analysis of the SARS virus Nsp15 endoribonuclease: Identification of residues affecting hexamer formation. J Mol Biol. 353, 1106–1117 (2005).

58. K. Bhardwaj, L. Guarino, C. C. Kao, The Severe Acute Respiratory Syndrome Coronavirus Nsp15 Protein Is an Endoribonuclease That Prefers Manganese as a Cofactor. J Virol. 78, 12218–12224 (2004).

59. K. A. Ivanov, T. Hertzig, M. Rozanov, S. Bayer, V. Thiel, A. E. Gorbalenya, J. Ziebuhr, Major genetic marker of nidoviruses encodes a replicative endoribonuclease. Proc Natl Acad Sci U S A. 101, 12694–12699 (2004).

60. Y. Kim, J. Wower, N. Maltseva, C. Chang, R. Jedrzejczak, M. Wilamowski, S. Kang, V. Nicolaescu, G. Randall, K. Michalska, A. Joachimiak, Tipiracil binds to uridine site and inhibits Nsp15 endoribonuclease NendoU from SARS-CoV-2. Commun Biol. 4, 1–11 (2021).

61. J. Ortiz-Alcantara, K. Bhardwaj, S. Palaninathan, M. Frieman, R. S. Baric, C. C. Kao, Small molecule inhibitors of the SARS-CoV Nsp15 endoribonuclease. Virus Adaptation and Treatment. 2, 125–133 (2010).

62. Y. Chen, G. Varani, Protein families and RNA recognition. FEBS Journal. 272 (2005), pp. 2088– 2097.

63. F. Mn, W. Im, K. Jm, B. Kj, D. Lb, B. Mj, S. Re, Flipped over U: structural basis for dsRNA cleavage by the SARS-CoV-2 endoribonuclease. Nucleic Acids Res. 1, 13–14 (2022).

64. K. Bhardwaj, S. Palaninathan, J. M. O. Alcantara, L. L. Yi, L. Guarino, J. C. Sacchettini, C. C. Kao, Structural and functional analyses of the severe acute respiratory syndrome coronavirus endoribonuclease Nsp15. Journal of Biological Chemistry. 283, 3655–3664 (2008).

65. E. M. Lynch, J. M. Kollman, B. A. Webb, Filament formation by metabolic enzymes—A new twist on regulation. Curr Opin Cell Biol. 66, 28–33 (2020).

66. A. Douangamath, D. Fearon, P. Gehrtz, T. Krojer, P. Lukacik, C. D. Owen, E. Resnick, C. Strain-Damerell, A. Aimon, P. Ábrányi-Balogh, J. Brandão-Neto, A. Carbery, G. Davison, A. Dias, T. D. Downes, L. Dunnett, M. Fairhead, J. D. Firth, S. P. Jones, A. Keeley, G. M. Keserü, H. F. Klein, M. P. Martin, M. E. M. Noble, P. O’Brien, A. Powell, R. N. Reddi, R. Skyner, M. Snee, M. J. Waring, C. Wild, N. London, F. von Delft, M. A. Walsh, Crystallographic and electrophilic fragment screening of the SARS-CoV-2 main protease. Nat Commun. 11, 1–11 (2020).

67. M. Schuller, G. J. Correy, S. Gahbauer, D. Fearon, T. Wu, R. E. Díaz, I. D. Young, L. C. Martins, D. H. Smith, U. Schulze-Gahmen, T. W. Owens, I. Deshpande, G. E. Merz, A. C. Thwin, J. T. Biel, J. K. Peters, M. Moritz, N. Herrera, H. T. Kratochvil, A. Aimon, J. M. Bennett, J. B. Neto, A. E. Cohen, A. Dias, A. Douangamath, L. Dunnett, O. Fedorov, M. P. Ferla, M. R. Fuchs, T. J. Gorrie-Stone, J. M. Holton, M. G. Johnson, T. Krojer, G. Meigs, A. J. Powell, J. G. M. Rack, V. L. Rangel, S. Russi, R. E. Skyner, C. A. Smith, A. S. Soares, J. L. Wierman, K. Zhu, P. O Brien, N. Jura, A. Ashworth, J. J. Irwin, M. C. Thompson, J. E. Gestwicki, F. von Delft, B. K. Shoichet, J. S. Fraser, I. Ahel, SARS-CoV-2 identified through crystallographic screening and computational docking. Sci Adv. 7, 25 (2021).

68. J. A. Newman, A. Douangamath, S. Yadzani, Y. Yosaatmadja, A. Aimon, J. Brandão-Neto, L. Dunnett, T. Gorrie-stone, R. Skyner, D. Fearon, M. Schapira, F. von Delft, O. Gileadi, Structure, mechanism and crystallographic fragment screening of the SARS-CoV-2 NSP13 helicase. Nature Communications 2021 12:1. 12, 1–11 (2021).

69. T. C. M. Consortium, H. Achdout, A. Aimon, E. Bar-David, H. Barr, A. Ben-Shmuel, J. Bennett, M. L. Boby, B. Borden, G. R. Bowman, J. Brun, S. Bvnbs, M. Calmiano, A. Carbery, E. Cattermole, E. Chernyshenko, J. D. Chodera, A. Clyde, J. E. Coffland, G. Cohen, J. Cole, A. Contini, L. Cox, M. Cvitkovic, A. Dias, K. Donckers, D. L. Dotson, A. Douangamath, S. Duberstein, T. Dudgeon, L. Dunnett, P. K. Eastman, N. Erez, C. J. Eyermann, M. Fairhead, G. Fate, D. Fearon, O. Fedorov, M. Ferla, R. S. Fernandes, L. Ferrins, R. Foster, H. Foster, R. Gabizon, A. Garcia-Sastre, V. O. Gawriljuk, P. Gehrtz, C. Gileadi, C. Giroud, W. G. Glass, R. Glen, I. Glinert, A. S. Godoy, M. Gorichko, T. Gorrie-Stone, E. J. Griffen, S. H. Hart, J. Heer, M. Henry, M. Hill, S. Horrell, M. F. D. Hurley, T. Israely, A. Jajack, E. Jnoff, D. Jochmans, T. John, S. de Jonghe, A. L. Kantsadi, P. W. Kenny, J. L. Kiappes, L. Koekemoer, B. Kovar, T. Krojer, A. A. Lee, B. A. Lefker, H. Levy, N. London, P. Lukacik, H. B. Macdonald, B. MacLean, T. R. Malla, T. Matviiuk, W. McCorkindale, B. L. McGovern, S. Melamed, O. Michurin, H. Mikolajek, B. F. Milne, A. Morris, G. M. Morris, M. J. Morwitzer, D. Moustakas, A. M. Nakamura, J. B. Neto, J. Neyts, L. Nguyen, G. D. Noske, V. Oleinikovas, G. Oliva, G. J. Overheul, D. Owen, V. Psenak, R. Pai, J. Pan, N. Paran, B. Perry, M. Pingle, J. Pinjari, B. Politi, A. Powell, R. Puni, V. L. Rangel, R. N. Reddi, S. P. Reid, E. Resnick, E. G. Ripka, M. C. Robinson, R. P. Robinson, J. Rodriguez-Guerra, R. Rosales, D. Rufa, C. Schofield, M. Shafeev, A. Shaikh, J. Shi, K. Shurrush, S. Singh, A. Sittner, R. Skyner, A. Smalley, M. D. Smilova, L. J. Solmesky, J. Spencer, C. Strain-Damerell, V. Swamy, H. Tamir, R. Tennant, W. Thompson, A. Thompson, W. Thompson, S. Tomasio, A. Tumber, I. Vakonakis, R. P. van Rij, L. Vangeel, F. S. Varghese, M. Vaschetto, E. B. Vitner, V. Voelz, A. Volkamer, F. von Delft, A. von Delft, M. Walsh, W. Ward, C. Weatherall, S. Weiss, K. M. White, C. F. Wild, M. Wittmann, N. Wright, Y. Yahalom-Ronen, D. Zaidmann, H. Zidane, N. Zitzmann, bioRxiv, in press, doi:10.1101/2020.10.29.339317.

