## Supplemental material 1 for "Allosteric regulation and crystallographic fragment screening of SARS-CoV-2 NSP15 endoribonuclease"

**Supplementary Material Information**

**Fig. S1.** A) Diagrams of different NendoU constructs. B) Gel filtration profile of NendoU$^{hex}$, showing peaks of hexamers (1), trimers (2) and monomers (3). Gel filtration profile of NendoU$^{mon}$, showing peaks of hexamers (4)

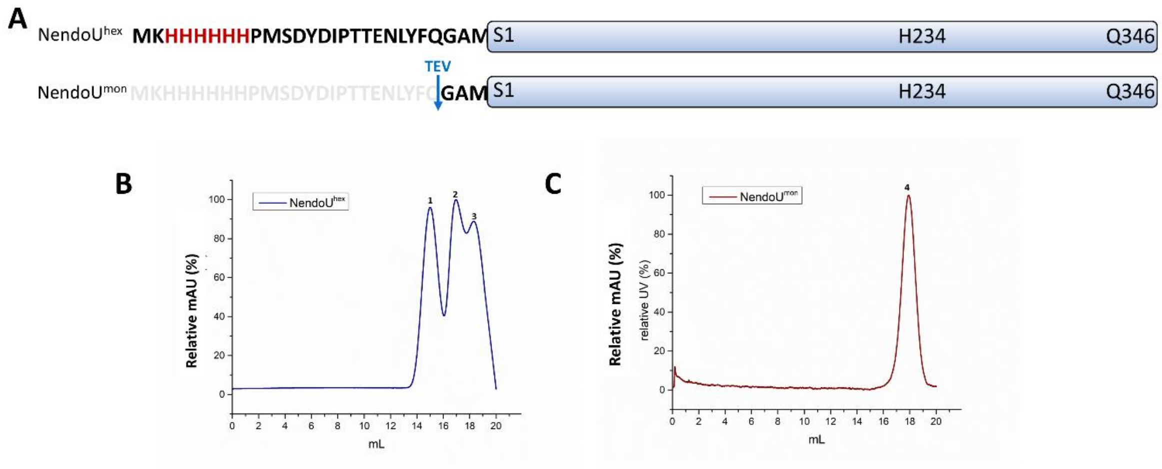


**Fig. S2.** Analytical Size exclusion chromatography profile of NendoU. A) calibration curve using standard proteins. B) Size exclusion chromatography profile of NendoU$^{hex}$, with a total mass of 204 kDa. C) Size exclusion chromatography profile of NendoU$^{mon}$, with a total mass of 34 kDa. In B and C, the sample containing peak is pointed with a black arrow.

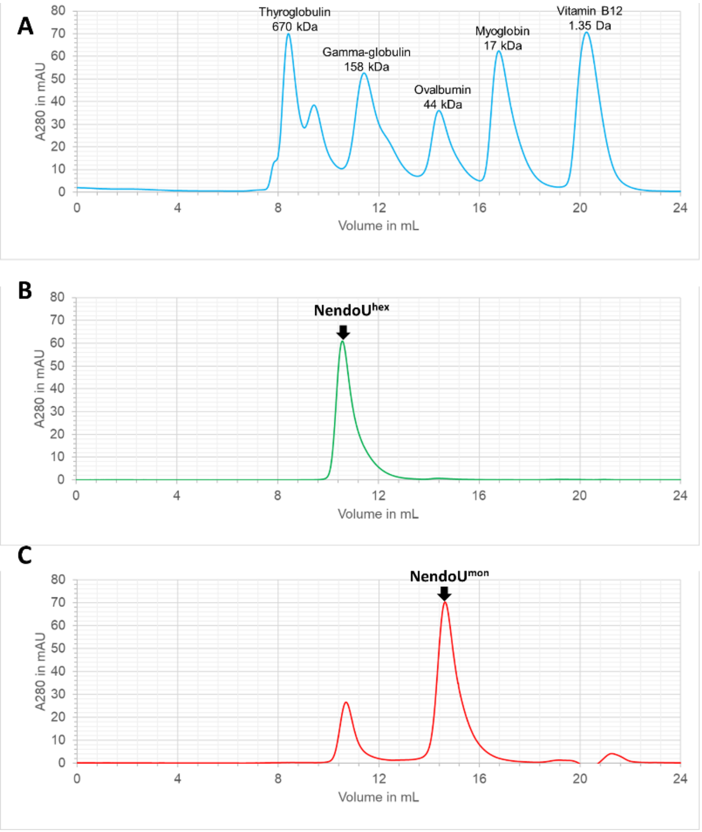


**Fig S3.** Mass spectroscopy profile of NendoU. In A), the mass spectra of full cleaved NendoU$^{mon}$. In B), the mass spectra of NendoU$^{hex}$ showing the mass of NSP15 with the additional expression N-terminal residues

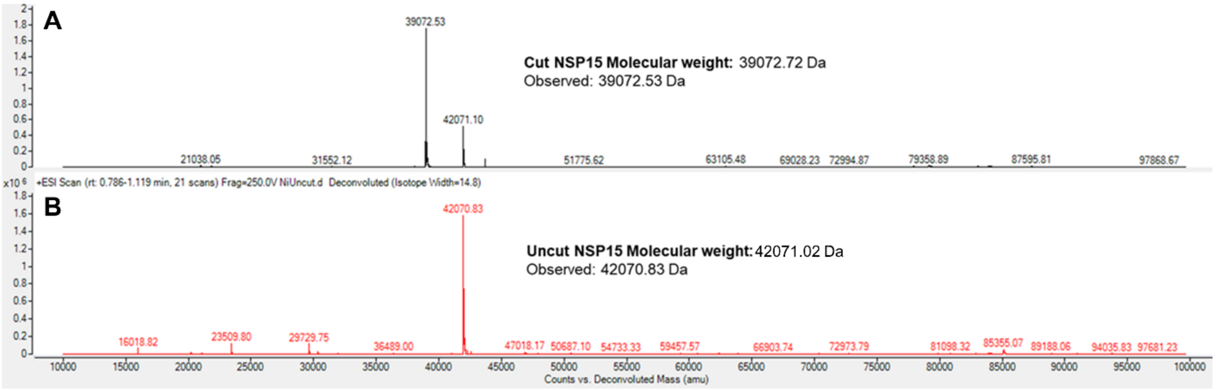

**Fig S4.** Native mass profiles of NendoU$^{mon}$ and NendoU$^{hex}$. A) Native mass spectra of NendoU$^{mon}$ sample, showing that the majority of the sample is presented as folded monomers. Is also possible to observe the presence of folded dimers, trimers and pentamers/hexamers B) Native mass spectra of NendoU$^{hex}$ sample, showing that the majority of the sample is presented as folded hexamers or folded monomers. Is also possible to observe the presence of folded dodecamers in the sample.

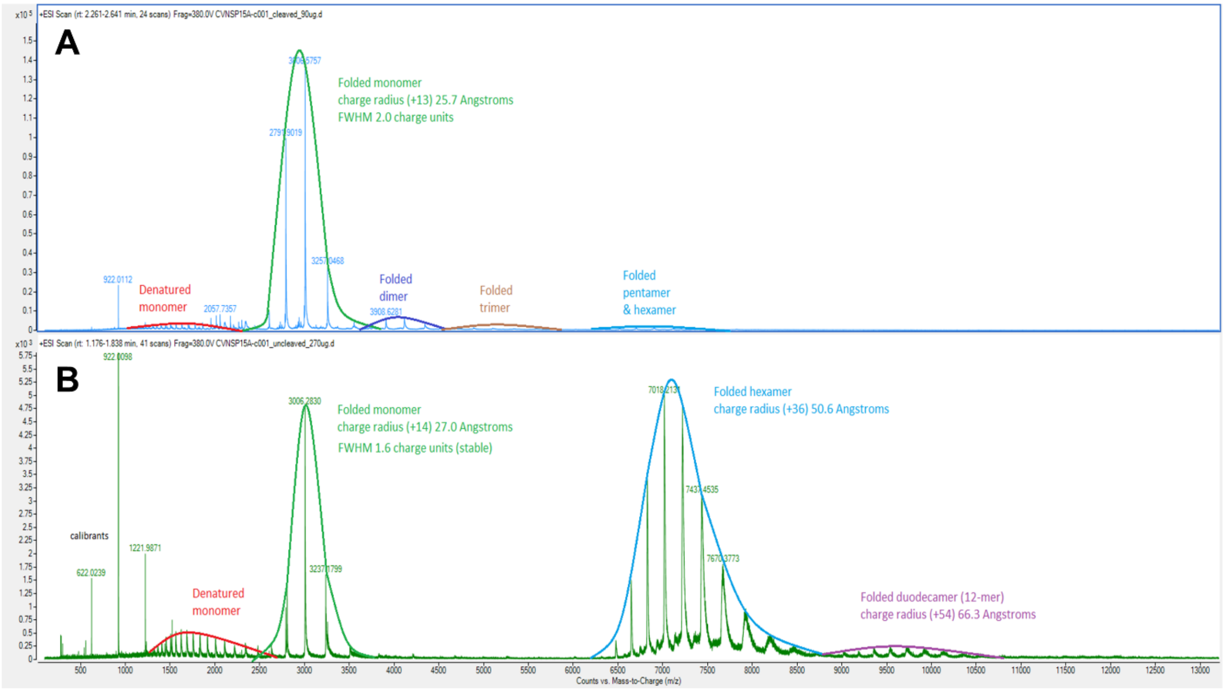

Fig S5. A) Crystal structure of the three domain NSP15 monomer. N-terminal domain is colored in shades of blue, middle domain is colored in shades of green and C-terminal catalytic domain is colored in shades of red-yellow. B) Front (left) and top (right) view of the NendoU hexamer. One of the chains is showed as cartoon (colored as A), while the other five are showed as spheres. C) Topology model of NSP15, following the same color pattern of A. Cylinders represent helices, with arrows are representing strands.

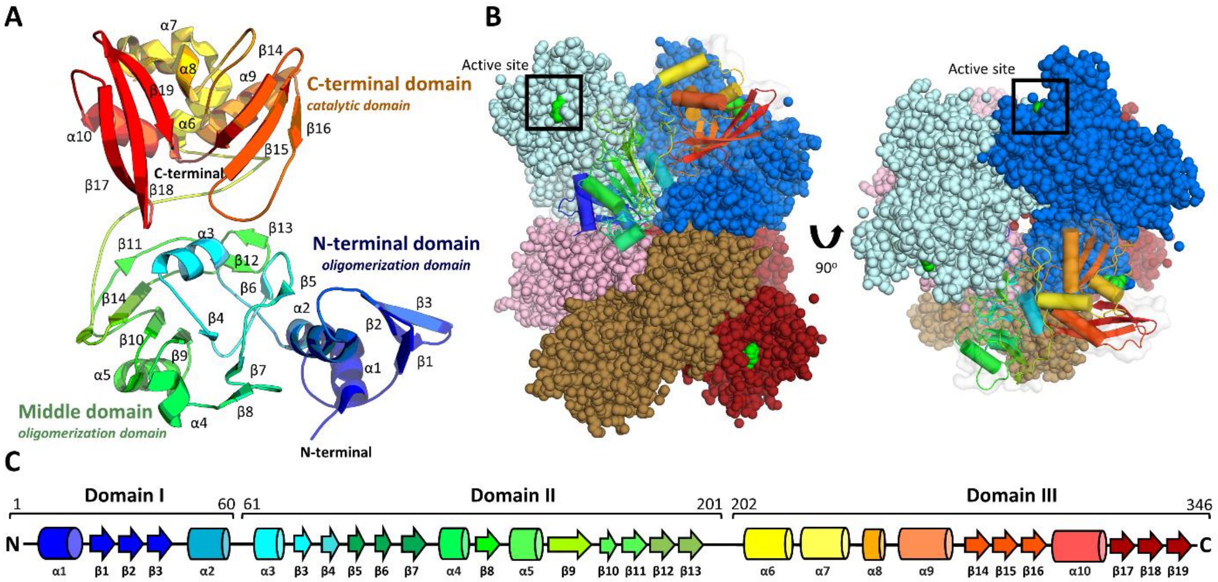

Fig. S6. Cryo-EM data processing schematic of NendoU$^{hex}$ collected in HEPES pH 7.5.

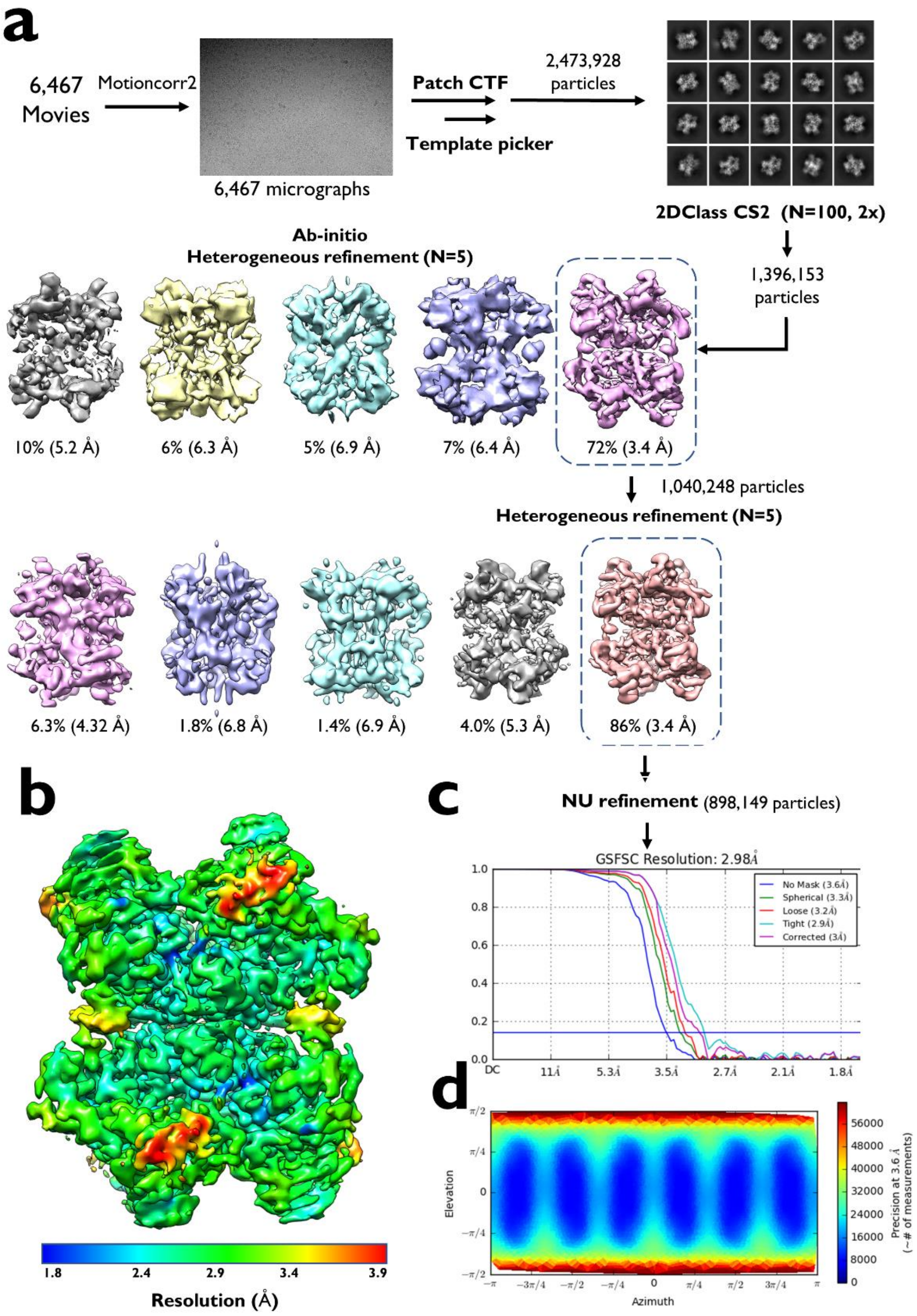

Fig. S7. Cryo-EM data processing schematic of NendoU$^{hex}$ collected in BIS-Tris pH 6.0.

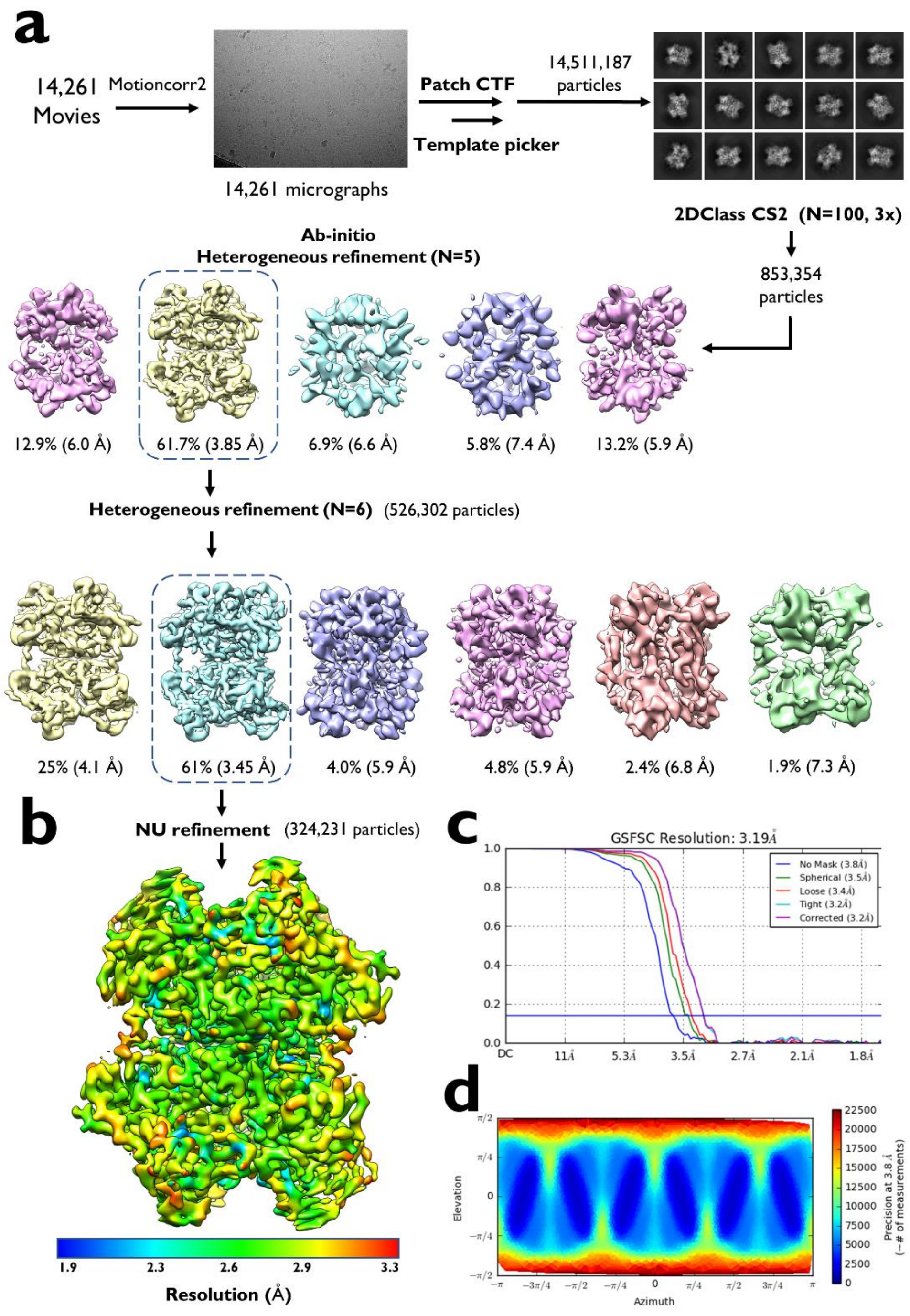

Fig. S8. Cryo-EM data processing schematic of NendoU$^{hex}$ collected in PBS pH 6.0.

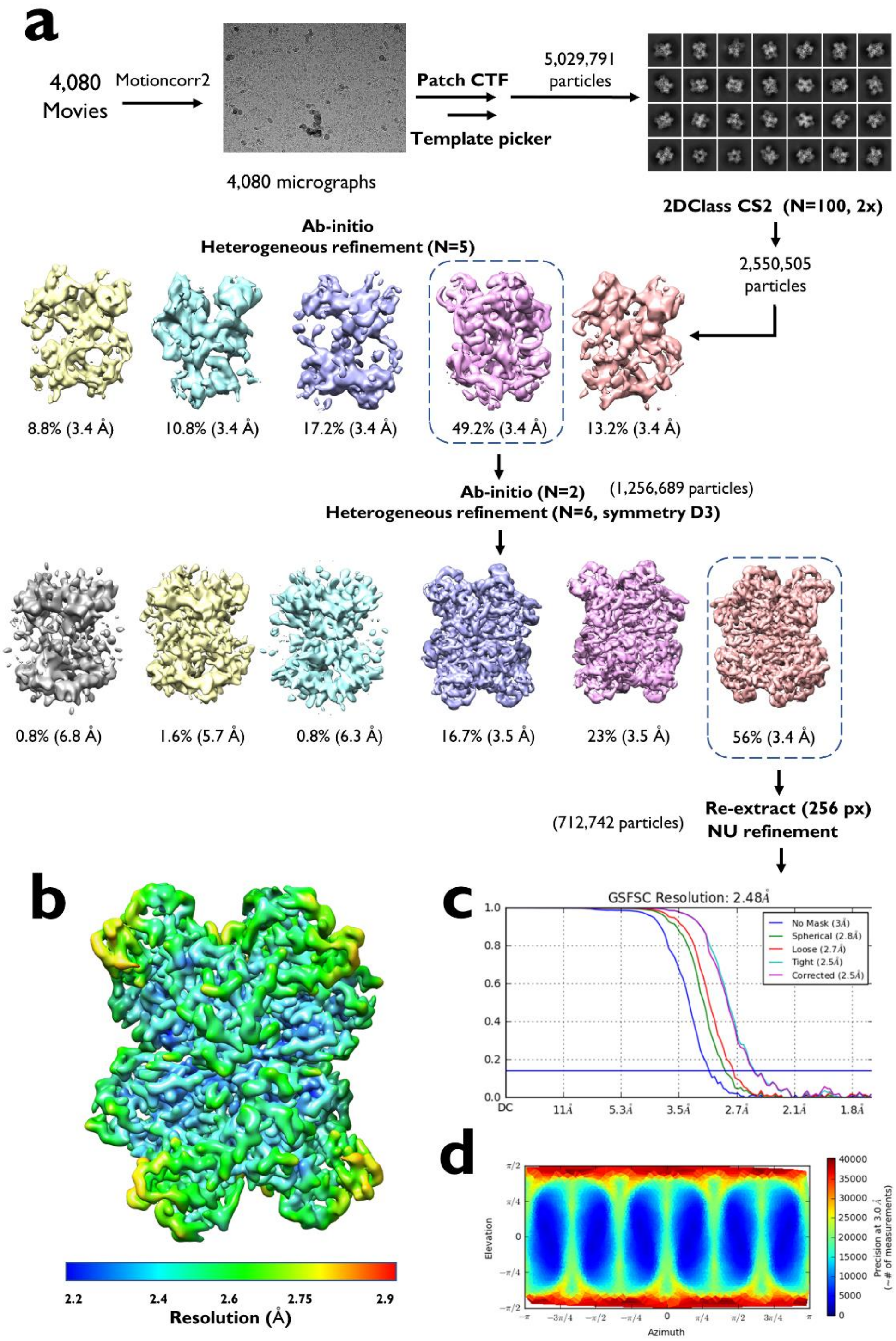

Fig. S9. A and B) Cryo-EM images of NendoU$^{hex}$ collected in HEPES pH 7.5.

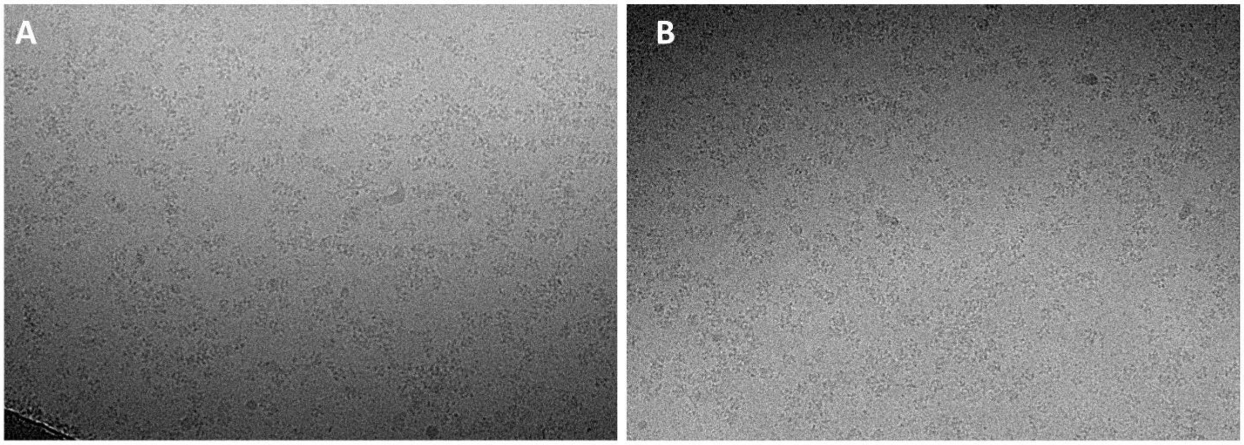


Fig. S10. A) Cryo-EM images of NendoU$^{hex}$ collected in Bis-tris pH 6.0. B) 2D classes of stacked particles. C) Low resolution volume reconstruction of stacked particles, with fitted model from PDB 7KF4.

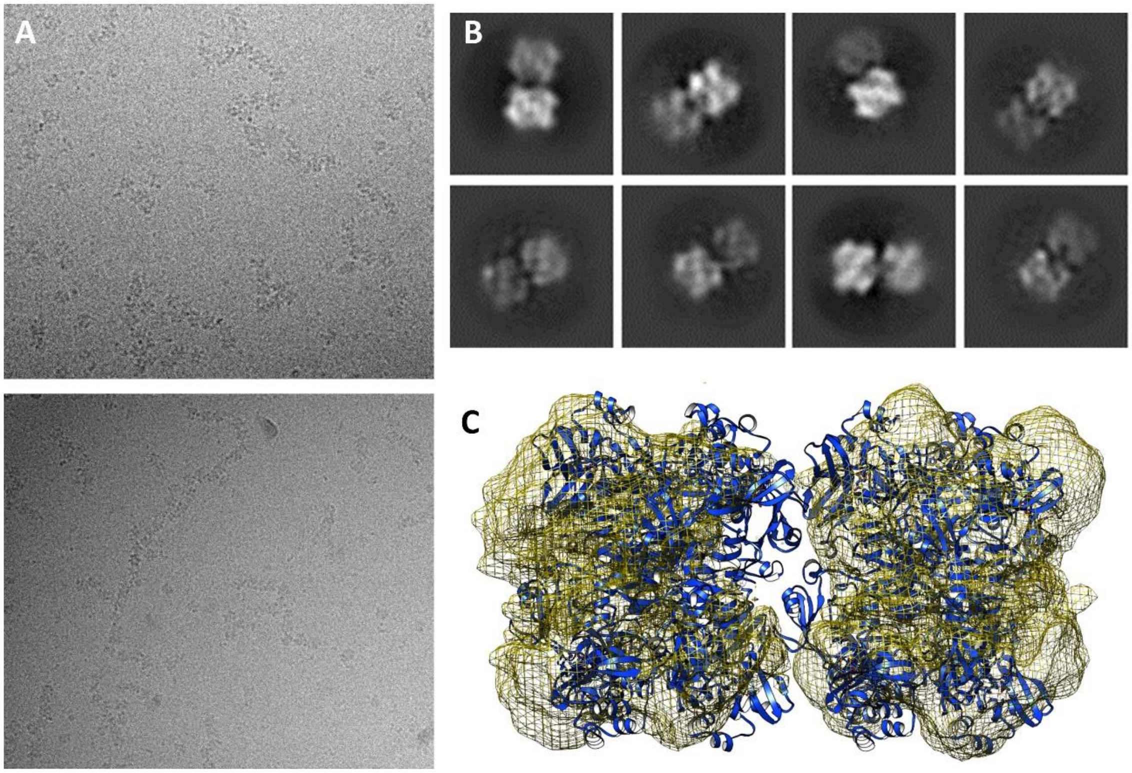

Fig S11. A) 2D crystals formed after adding DNA oligoDT to the sample. B) 2D classes obtained from 2D crystals samples. C) Fourier transformation of 2D crystals images.

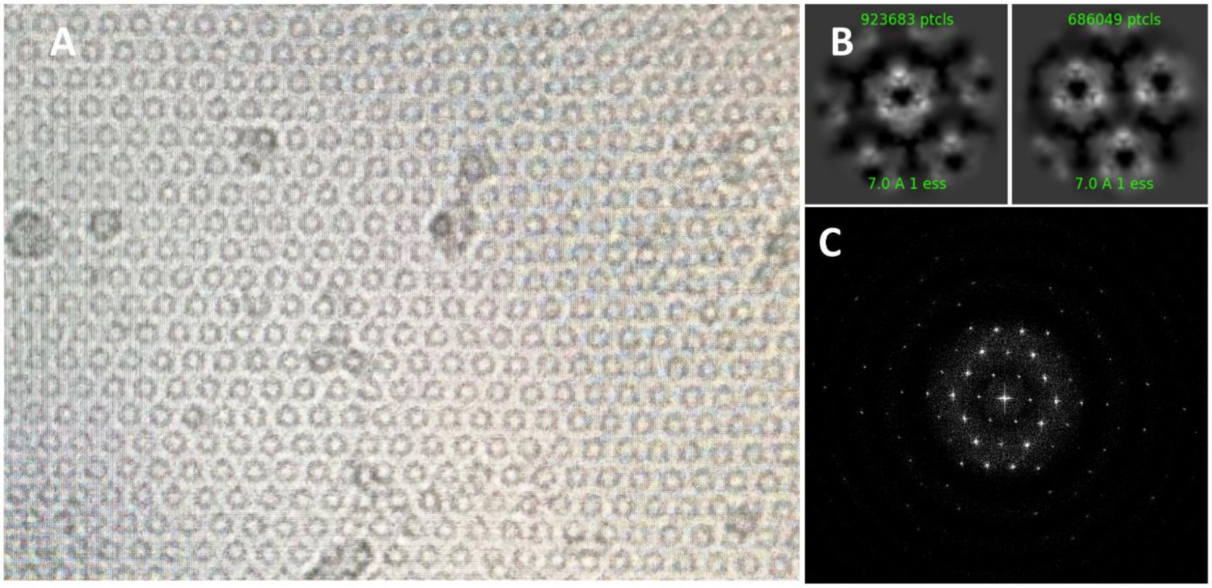

Fig. S12. Different crystals packing between crystals obtained in presence or absence of DNA. A) Crystal packing from crystals obtained without the presence of DNA. B) Crystal packing from crystals obtained in the presence of DNA oligoDT$_{30}$.

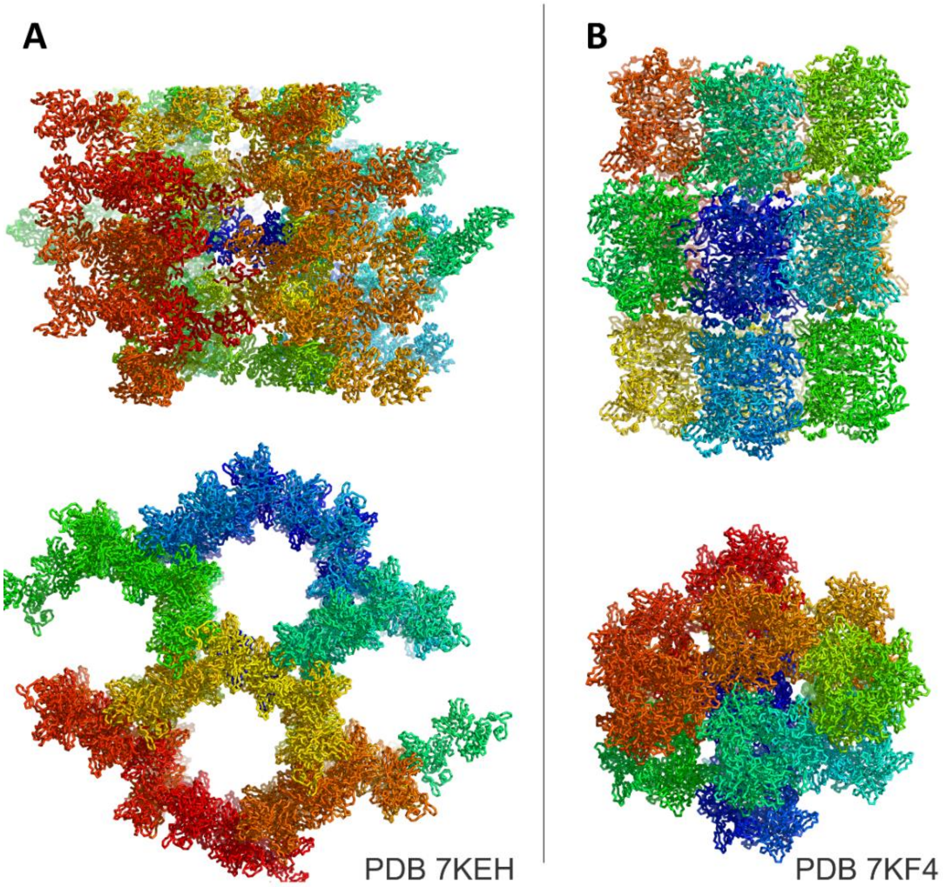


Fig S13. A) Structural comparison between thymidine, uridine, FUZS-5 and LIZA-7 fragments. B) Effect of different FUZS-5 concentration in NendoU relative enzymatic activity determined using fluorogenic substrate.

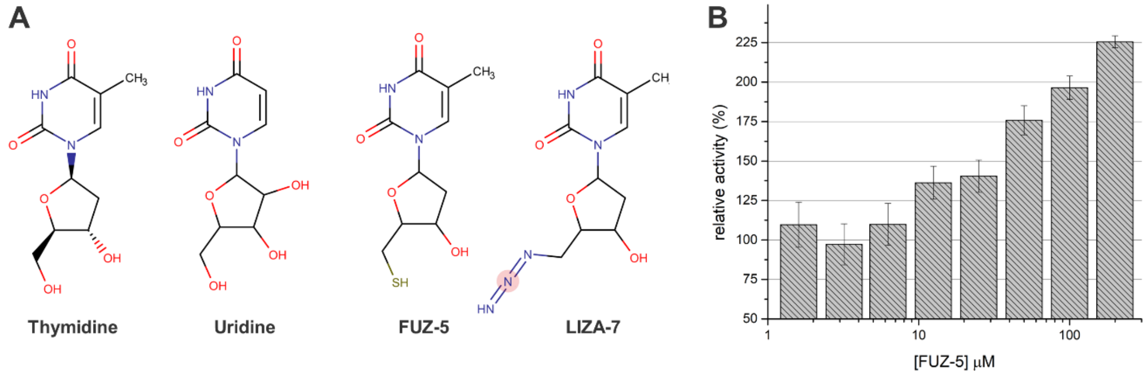

Fig S14. A) Distinct binding modes of NendoU to dsRNA, generated expanding symmetry of PDB 7TJ2. B) Sequence of nsp15 colored according to conservation. C) Surface view of nsp15 structure colored according to conservation. Conservation analyses were performed with Consurf.

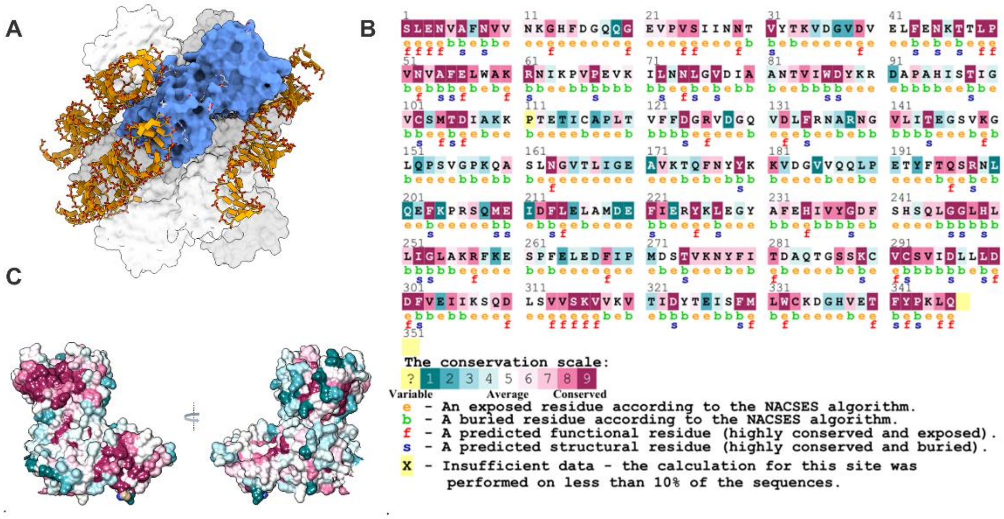

Table S1. Activity profile of NendoU$^{mon}$ and NendoU$^{hex}$ in different buffers.

| | | | | | **NendoU$^{mon}$** | | **NendoU$^{hex}$** | |
|---|---|---|---|---|---|---|---|---|
| **Buffer** | **Buffer concentration (mM)** | **NaCL (mM)** | **Glycerol (%)** | **pH** | **$v_o$ (RFU/min)** | **Error** | **$v_o$ (RFU/min)** | **Error** |
| Sodium acetate | 50 | 150 | 0 | 5.0 | 19.57 | 0.126 | 241.09 | 18.78 |
| Sodium cacodylate (HCl) | 50 | 150 | 0 | 6.0 | 15.49 | 0.078 | 998.43 | 31.25 |
| MES (NaOH) | 50 | 150 | 0 | 6.0 | 16.55 | 0.067 | 92.58 | 2.58 |
| BIS-TRIS (NaOH) | 50 | 150 | 0 | 6.5 | 15.62 | 0.14 | 902.76 | 13.90 |
| Imidazole (HCl) | 50 | 150 | 0 | 7.0 | 2.84 | 0.15 | 804.66 | 3.45 |
| MOPS (NaOH) | 50 | 150 | 0 | 7.2 | 1.42 | 0.09 | 100.27 | 1.62 |
| Tricine (NaOH) | 50 | 150 | 0 | 8.0 | 3.49 | 0.14 | 63.67 | 7.15 |
| TRIS (HCl) | 50 | 150 | 0 | 8.0 | - | - | 174.53 | 5.35 |
| TRIS (HCl) | 50 | 150 | 0 | 8.5 | 2.76 | 0.17 | 43.06 | 2.10 |
| Bicine (NaOH) | 50 | 150 | 0 | 8.3 | 6.00 | 0.24 | - | - |
| HEPES (NaOH) | 50 | 150 | 0 | 7.5 | - | - | 29.74 | 0.81 |
| Glycine (NaOH) | 50 | 150 | 0 | 9.0 | 0.90 | 0.142 | - | - |
| CHES (NaOH) | 50 | 150 | 0 | 9.5 | - | - | 27.56 | 1.36 |
| Sodium borate | 50 | 150 | 0 | 10.0 | - | - | - | - |
| HEPES (NaOH) | 50 | 0 | 0 | 7.5 | - | - | 156.54 | 2.9 |
| HEPES (NaOH) | 50 | 50 | 0 | 7.5 | - | - | 83.65 | 1.20 |
| HEPES (NaOH) | 50 | 500 | 0 | 7.5 | 0.33 | 0.0 | 46.08 | 2.65 |
| HEPES (NaOH) | 50 | 150 | 5 | 7.5 | - | - | 73.07 | 5.03 |
| HEPES (NaOH) | 50 | 150 | 10 | 7.5 | 0.012 | 0.037 | 136.80 | 3.42 |
| PBS | - | - | - | 7.4 | - | - | 74.19 | 2.97 |

Table S2. Data collection and processing statistics for cryo-EM models of NendoU

| | TRIS-HCl pH 7.5 | BIS-TRIS pH 6.0 | PBS pH 6.0 |
|---|---|---|---|
| **Data collection** | | | |
| C2 (µm) | 50 | 70 | 70 |
| Spot size | 6 | 6 | 5 |
| Illuminiation (µm) | 1.1 | 0.99 | 1.25 |
| Magnification | 105k | 105k | 105k |
| Camera | K3 super resolution | K3 super resolution | K3 super resolution |
| Slit width | 20 eV | 20 eV | 20 eV |
| Pixel (physical) (Å) | 0.831 | 0.831 | 0.831 |
| Dose rate (e/px/s) | 14.5 | 15.01 | 18 |
| exposure time (s) | 2.02 | 1.99 | 1.6 |
| total dose (e/Å2) | 42.41 | 43.25 | 41.7 |
| fractions | 40 | 40 | 40 |
| dose per frame (e/Å2/frame) | 1.06 | 1.08 | 1.04 |
| Movies collected | 6,467 | 14,261 | 4,080 |
| **Model Refinement** | | | |
| Chains | 6 | 6 | 6 |
| Atoms | 16350 | 16386 | 16416 |
| Water | 0 | 0 | 0 |
| Bonds (RMSD) | | | |
| Length (Å) | 0.009 (0) | 0.008 (0) | 0.006 (0) |
| Angles (°) | 0.756 (1) | 0.695 (3) | 0.553 (1) |
| MolProbity score | 1.96 | 1.71 | 1.85 |
| Clash score | 12.75 | 6.92 | 5.72 |
| Ramachandran plot (%) | | | |
| Outliers | 0 | 0 | 0 |
| Allowed | 4.88 | 4.73 | 2.03 |
| Favored | 95.12 | 95.27 | 97.97 |
| Rama-Z (Ramachandran plot Z-score, RMSD) | | | |
| whole (N = 2076) | 1.42 (0.18) | 0.98 (0.19) | 0.22 (0.17) |
| helix (N = 516) | 1.13 (0.20) | 0.83 (0.22) | 0.20 (0.22) |
| sheet (N = 312) | 0.81 (0.28) | 0.54 (0.32) | 0.93 (0.30) |
| loop (N = 1248) | 0.83 (0.18) | 0.52 (0.18) | 0.56 (0.16) |
| Rotamer outliers (%) | 0 | 0 | 4.96 |
| Cβ outliers (%) | 0 | 0 | 0 |
| Peptide plane (%) | | | |
| Cis proline/general | 0.0/0.0 | 0.0/0.0 | 0.0/0.0 |
| Twisted proline/general | 0.0/0.0 | 0.0/0.0 | 0.0/0.0 |
| CaBLAM outliers (%) | 1.55 | 1.60 | 1.02 |
| d FSC model (0/0.143/0.5) | 2.9/3.0/3.3 | 3.1/3.2/3.4 | 2.4/2.5/2.7 |
| Model vs. Data | | | |
| CC (mask) | 0.75 | 0.81 | 0.83 |
| CC (box) | 0.75 | 0.77 | 0.82 |
| CC (peaks) | 0.7 | 0.74 | 0.79 |
| CC (volume) | 0.74 | 0.79 | 0.82 |
| PDB deposition code | 7RB0 | 7RB2 | 7ME0 |
| EMDB deposition code | EMD-24391 | EMD-24392 | EMD-23786 |

Table S3. Data collection and processing statistics for X-ray models of NendoU

| | Dihedral AU | Hexamer AU | Dihedral AU |
|---|---|---|---|
| Beamline | MANACA | MAXIV | MAXIV |
| Wavelenght (Å) | 1.335 | 0.976 | 0.976 |
| Resolution range (Å) | 45.0 - 2.9 (3.0 - 2.9) | 49.16 - 2.61 (2.7 - 2.6) | 84.6 - 2.6 (2.74 - 2.6) |
| Space group | P 63 | P 2 21 21 | P 63 |
| Unit cell (a, b & c; Å, angles) | 150.6 150.6 110.2 90 90 120 | 85.1 151.1 199.2 90 90 90 | 150.9 150.9 111.1 90 90 120 |
| Unique Reflections | 31369 (3117) | 78886 (7756) | 37814 (1889) |
| Multiplicity | 10.0 (10.2) | 13.8 (13.9) | 29.8 (30.1) |
| Completeness (%) | 99.04 (98.67) | 99.73 (99.73) | 94.8 (60.3) |
| Mean I/sigma(I) | 5.0 (1.0) | 8.24 (0.72) | 7.6 (1.6) |
| Rpim (%) | 0.188 (0.912) | 0.06367 (0.9967) | 0.19 (0.60) |
| CC1/2 | 0.96 (0.31) | 0.998 (0.527) | 0.99 (0.37) |
| *Rwork* | 0.1949 (0.3064) | 0.2331 (0.3987) | 0.19 |
| *Rfree* | 0.2332 (0.3290) | 0.2573 (0.4120) | 0.22 |
| Number of atoms | | | |
| ligands | 10 | 78 | 48 |
| waters | 357 | 429 | 468 |
| Protein residues | 696 | 2088 | 696 |
| RMS(bonds) (Å) | 0 | 14 | 13 |
| RMS(angles) (o) | 1,78 | 1.69 | 1.76 |
| Ramachandran favored (%) | 96.68 | 97.50 | 97.25 |
| Ramachandran outliers (%) | 0 | 0.10 | 0 |
| Clashscore | 2 | 2.58 | 2.88 |
| Average B-factors (Å²) | | | |
| Macromolecules | 54.18 | 74.33 | 58.10 |
| Ligands | 89.87 | 108.38 | 67.35 |
| Solvent | 43.54 | 57.70 | 53.28 |
| PDB code | 7KEG | 7KF4 | 7KEH |

NMR spectra for **5’-S-Acetyl-5’-deoxy-5’-thiothymidine**

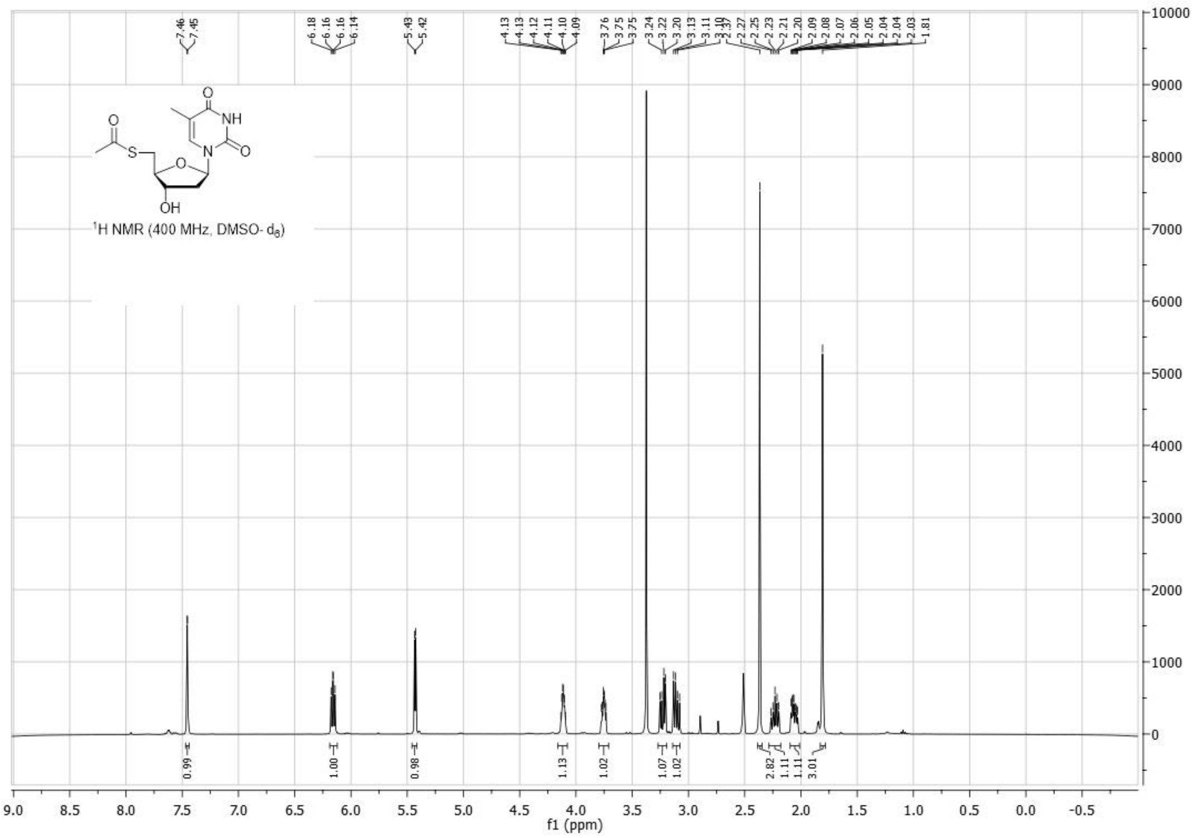


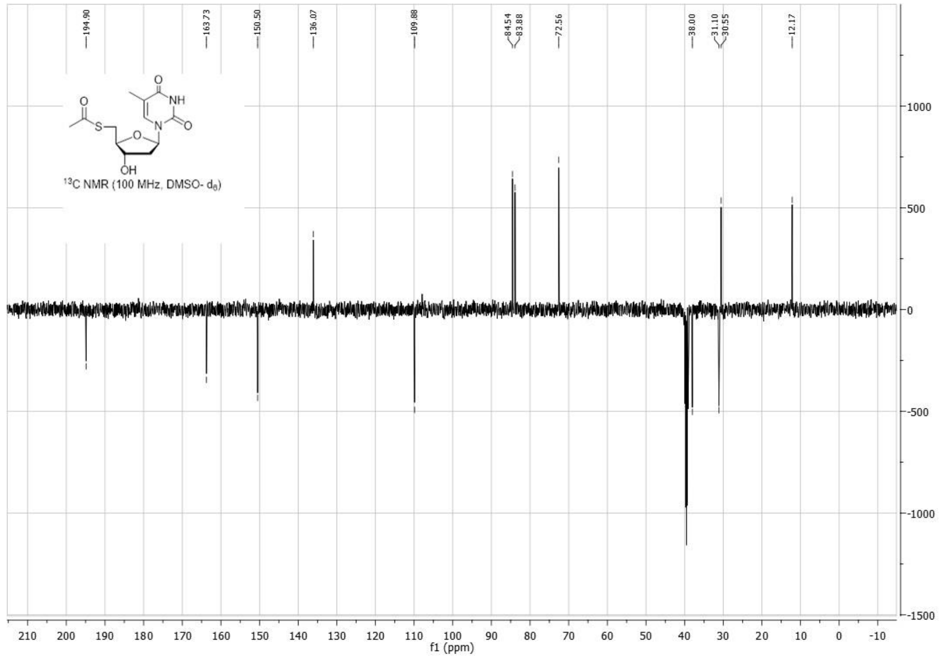

NMR spectra for **5'-deoxy-5'-thiothymidine**

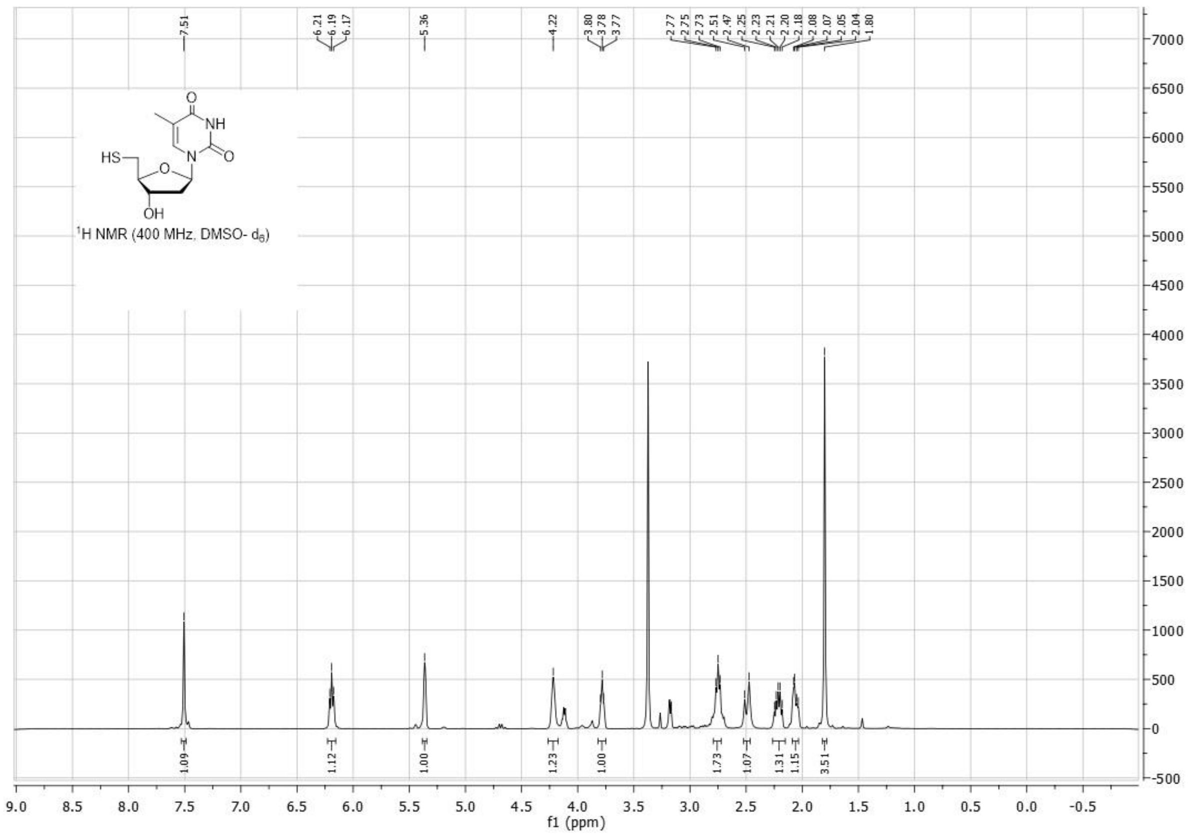


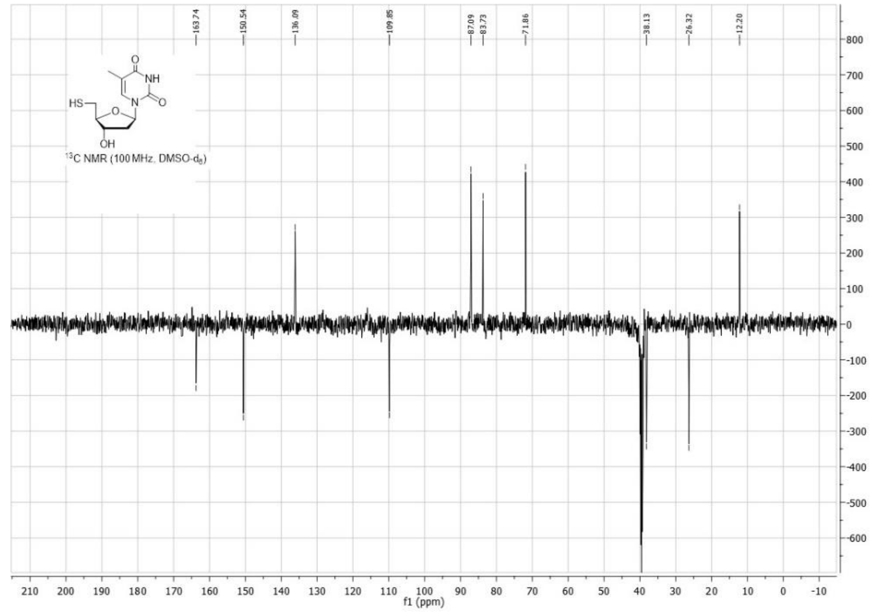
